# Arousal modulates functional connectivity through structured and hemispherically asymmetric community architecture during wakefulness

**DOI:** 10.64898/2026.01.06.697875

**Authors:** Xiangyu Kong, Siyu Li, Gaolang Gong

## Abstract

Arousal fluctuates continuously during wakefulness, yet how these moment-to-moment variations shape large-scale functional connectivity (FC) remains unclear. Here, we combined 7T fMRI with concurrent pupillometry to quantify, for every functional connection, how time-varying FC covaries with spontaneous arousal in the awake human brain. Rather than exerting a uniform influence across the connectome, arousal organized FC into a low-dimensional set of seven connectivity communities, each defined by characteristic network compositions. These communities exhibited systematic hemispheric asymmetries, specifically identifying a “left-hemisphere centripetal architecture” where the left hemisphere serves as a structural sink for the asymmetric convergence of arousal-modulated signals. Importantly, hemispheric asymmetry did not arise from global shifts in connectivity strength, but instead reflected structured spatial heterogeneity embedded within community architecture. This modular and asymmetric organization was highly preserved during naturalistic movie watching, indicating that arousal-related modulation of FC reflects intrinsic principles that generalize across awake cognitive contexts. Together, these findings demonstrate that moment-to-moment arousal fluctuations shape large-scale FC through structured, hemispherically asymmetric network organization during wakefulness.

## Introduction

Arousal is a core dimension of brain state that shapes perception, attention, and the coordination of large-scale neural systems. Across major transitions—such as wakefulness to sleep, anesthesia, or disorders of consciousness—substantial reorganization of functional connectivity (FC) has been consistently observed (Chow et al., 2013; Tagliazucchi et al., 2016; Demertzi et al., 2019; Banks et al., 2020; Damaraju et al., 2020; Huang et al., 2020; Jang et al., 2024). These state-based findings demonstrate that arousal fundamentally influences whole-brain communication patterns. However, such approaches typically contrast discrete arousal states and therefore provide limited insight into how moment-to-moment fluctuations in arousal within the awake brain influence the organization of functional connectivity.

Increasing evidence shows that arousal varies continuously even during stable wakefulness, including during resting-state fMRI and ongoing cognitive engagement. These spontaneous fluctuations, often indexed by pupil diameter (Reimer et al., 2014; McGinley et al., 2015; Joshi & Gold, 2020), modulate neural gain, sensory responses, and behavioral performance. Yet despite their ubiquity and behavioral relevance, it remains largely unknown how fine-grained arousal variations are expressed across the functional connectome. Prior work has primarily examined regional signal amplitude or isolated networks (Yellin et al., 2015; Schneider et al., 2016; Breeden et al., 2017; Podvalny et al., 2021; Sobczak et al., 2021; Lloyd et al., 2023), thereby leaving unresolved the fundamental question of whether the awake brain’s FC is uniformly sensitive to arousal or whether arousal instead imprints structured spatial patterns across functional networks.

A parallel question is whether arousal-modulated connectivity patterns show hemispheric asymmetry, given longstanding evidence for lateralized arousal, vigilance, and alerting mechanisms. Studies of unihemispheric sleep and lateralized arousal dynamics in animals (Rattenborg et al., 2000; Lyamin et al., 2016; Mascetti, Gian Gastone, 2016; Reicher et al., 2021; Fenk et al., 2023; Libourel et al., 2023), asymmetries in human EEG-based vigilance (Tamaki et al., 2016), and right-lateralized alerting functions in attention (Heilman & Abell, 1980; Sturm & Willmes, 2001; Shulman et al., 2010; Corbetta & Shulman, 2011) all suggest that arousal may modulate left and right hemispheric systems differently. Nevertheless, these observations have yet to be linked to the organization of whole-brain functional interactions, and it remains unknown whether such asymmetries manifest at the level of large-scale FC, and whether they reflect organized connectivity patterns rather than non-specific global effects.

To address these questions, we combined high-field fMRI with concurrent pupillometry to quantify, for every functional connection, how its connectivity covaries with spontaneous arousal fluctuations during wakefulness. This edgewise measure of arousal–time varying functional connectivity (tvFC) coupling provides a comprehensive map of where in the connectome arousal leaves its strongest imprint, without imposing predefined states or regional assumptions. Using this framework, we first test whether arousal sensitivity is spatially homogeneous or segregates into distinct sets of connections with similar coupling profiles. We next assess whether these spatially organized arousal-modulated patterns show systematic hemispheric asymmetry, with particular emphasis on attentional systems that show known lateralization. Finally, we evaluate the cross-context stability of this organizational structure by comparing resting state and naturalistic movie watching in the same participants. Together, these analyses delineate how moment-to-moment arousal fluctuations shape large-scale functional architecture in the awake human brain.

## Results

The processing procedure of estimating arousal–tvFC coupling from fMRI and pupillometry was illustrated in Figure 1. Here, we use the term arousal–tvFC coupling to refer to the regression-based estimate of how spontaneous arousal fluctuations modulate each functional connection over time.

**Figure 1.**
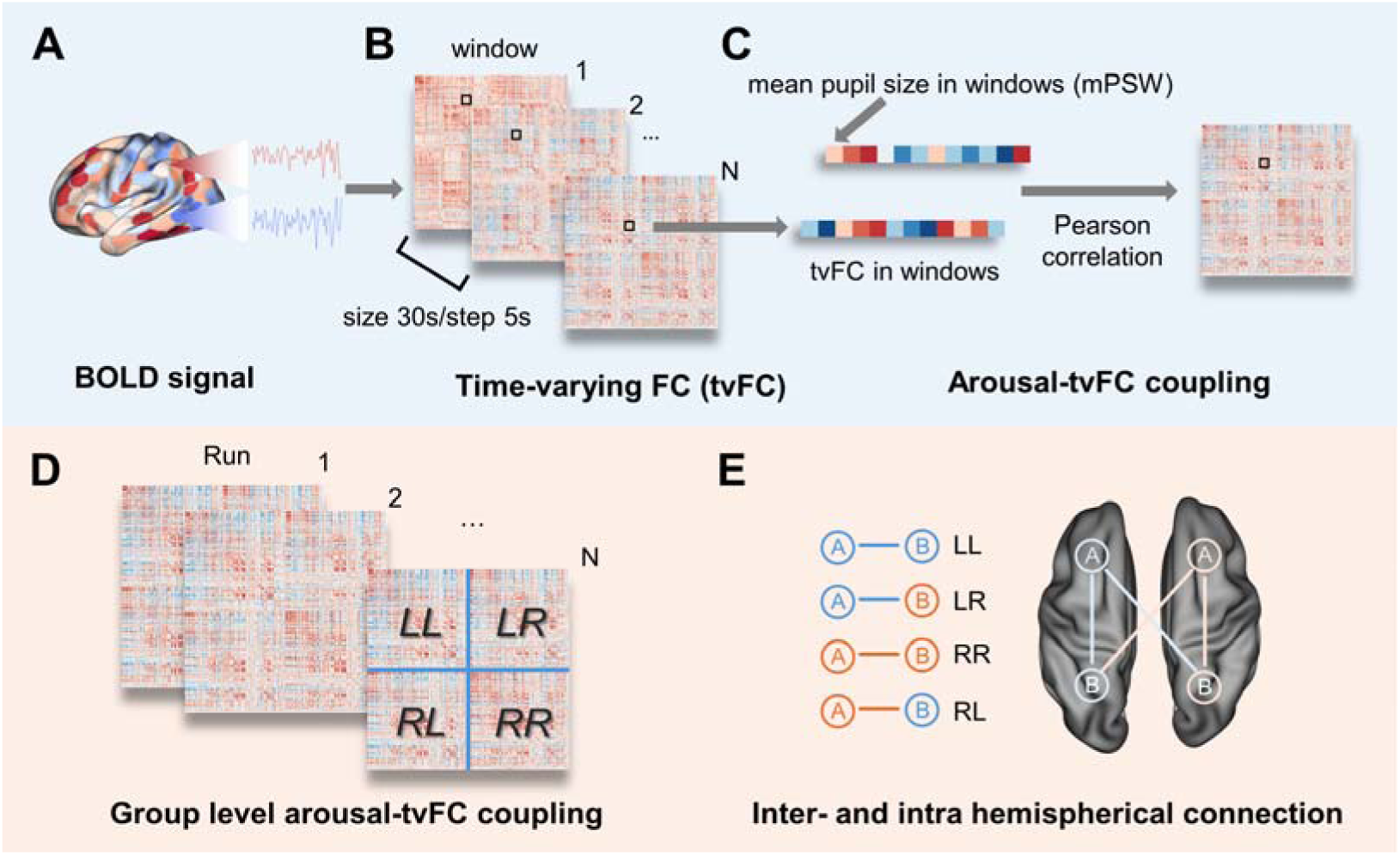
Pipeline for estimating arousal–tvFC coupling from fMRI and pupillometry. **(A)** Concurrent 7T fMRI and eye tracking were collected during resting state and naturalistic movie watching. **(B)** tvFC was computed over sliding windows for each pair of brain regions. **(C)** Pupil diameter was preprocessed to obtain a continuous arousal time series. Arousal–tvFC coupling for each connection was then defined as the Pearson correlation between its windowed FC time course and the corresponding arousal fluctuations. **(D)** These edgewise coupling values yielded a dense arousal–tvFC coupling matrix per run, which served as the input for analyses of community structure, hemispheric asymmetry, and cross-paradigm consistency. **(E)** The connections were categorized based on the hemispheres of the connected regions: Left-Left (LL) and Right-Right (RR) represent intra-hemispheric connections; Left-Right (LR) and Right-Left (RL) represent inter-hemispheric connections. This classification enabled hemisphere-specific analyses of arousal-tvFC coupling.

While our primary inferential analyses were conducted at the run level to leverage the high-density sampling of the HCP 7T dataset, we further validated the robustness of these findings using participant-level statistical summaries and resampling to account for within-participant dependencies (see Figure. S1-S2 in Supplementary Material).

### Arousal–tvFC coupling reveals seven distinct connectivity communities

For each functional connection, we quantified how strongly its time-varying connectivity covaried with moment-to-moment arousal, producing an edgewise arousal–tvFC coupling matrix per run. Across all participants and runs, these coupling profiles showed clear structure: connections did not exhibit uniform arousal sensitivity but instead formed separable groups.

Unsupervised clustering of edgewise coupling patterns identified seven stable connectivity communities (Fig. 2A). This solution was consistently favored across a broad range of cluster numbers, indicating that arousal–tvFC coupling is inherently low-dimensional. The seven communities captured distinct patterns of arousal sensitivity across the connectome. Projecting each community into canonical network pairs space (Thomas Yeo et al., 2011) showed that these communities were not random mixtures of connections. Instead, each displayed a characteristic distribution across network pairs (Fig. 2B). Some communities were enriched in network pairs linking heteromodal and unimodal systems (Mesulam, 1998), while others were dominated by heteromodal–heteromodal (H-H) or unimodal–unimodal (U-U) network pairs. Community participation entropy further highlighted this structure: unimodal–unimodal connections showed low entropy, indicating a restricted participation in a few communities, whereas H-H and heteromodal–unimodal (H-U) connections showed significantly higher entropy, indicating more diverse engagement across communities (F(2,88)=12.24, p = 4.38×10^-6^).

**Figure 2.**
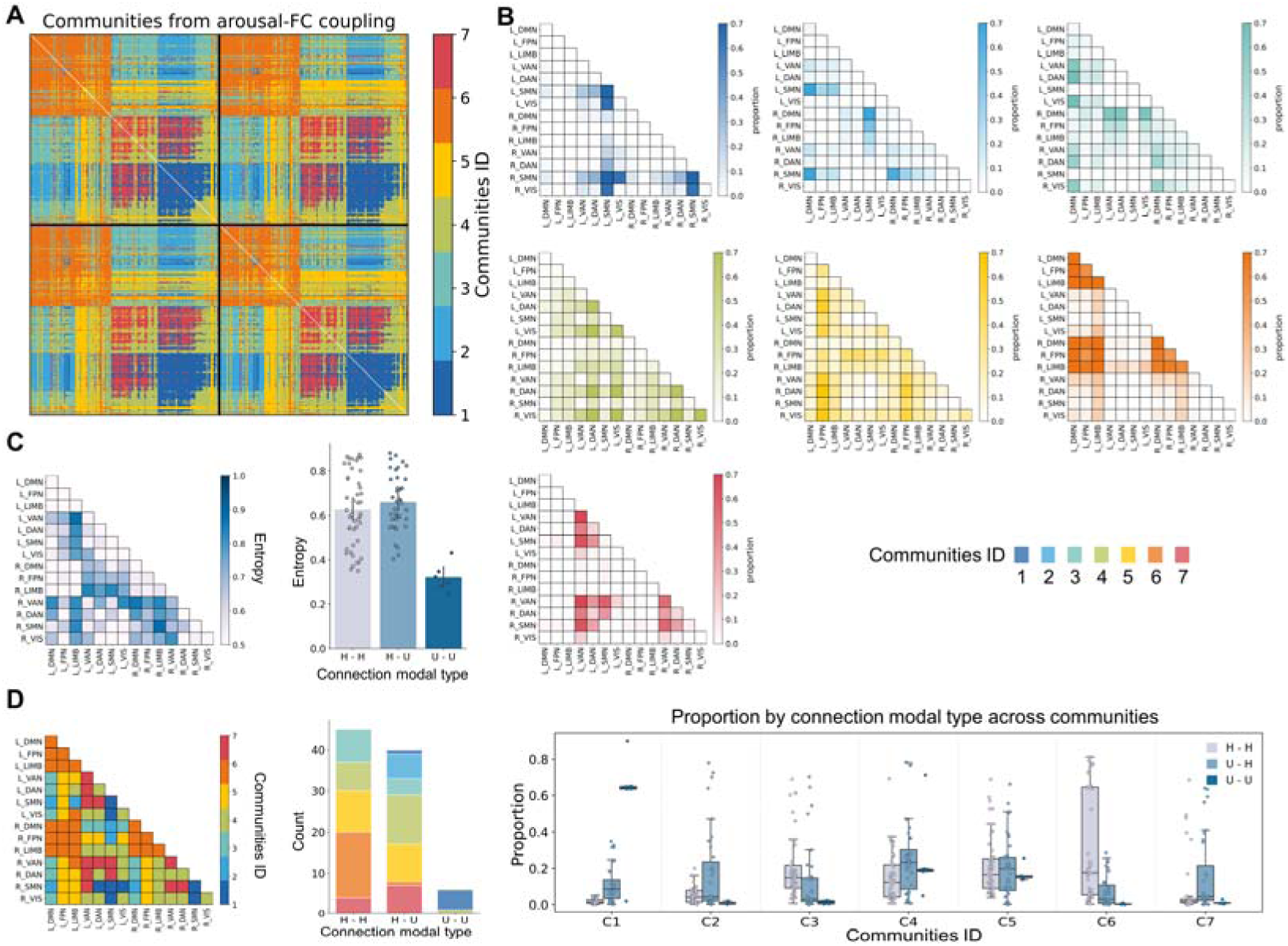
Arousal–tvFC coupling partitions the connectome into seven distinct connectivity communities. **(A)** Unsupervised clustering of edgewise arousal–tvFC coupling identified seven stable communities, demonstrating that arousal-linked modulation is organized into low-dimensional structure rather than uniformly distributed across connections. **(B)** Mapping communities onto network-pair space revealed distinct and reproducible composition profiles (upper panel), with some communities dominated by heteromodal interactions and others enriched in H–U or U–U network pairs (bottom panel). **(C)** Community participation entropy varied systematically across connection modal types: U–U network pairs showed lower entropy, indicating concentrated engagement in a small subset of communities, whereas H-H and H–U network pairs exhibited significantly higher entropy (F(2,88)=12.24, p = 4.38×10^-6^), reflecting broader distribution across communities. **(D)** Dominant-community assignments confirmed this organization, showing that heteromodal interactions load onto multiple arousal-sensitive communities, whereas U–U network pairs show more restricted community involvement.

Together, these results demonstrate that arousal does not uniformly modulate the connectome but instead engages a small number of organized connectivity communities, each with distinct network-level compositions.

The robustness of the seven-community architecture was cross-validated using both split-half and participant-level resampling strategies. As shown in **Figure S1**, both approaches yielded high alignment accuracy and consistently high Dice coefficients across all communities, confirming that the identified clusters are not artifacts of specific data partitioning.

### Arousal-modulated community architecture exhibit systematic hemispheric asymmetry

We next asked whether these arousal-modulated communities express hemispheric biases. Using integration and segregation indices derived from LL, RR, LR, and RL edge categories, we quantified, for each community, whether arousal preferentially modulated intra- versus inter-hemispheric connectivity and whether these effects favored one hemisphere.

At the network-pair level, several pairs showed significant lateralization compared with a spatial permutation null model (Fig. 3A–B). Importantly, lateralization was not global: only specific network pairs within communities exhibited robust hemispheric biases, while others remained symmetric. Some communities showed rightward integration, indicating stronger arousal-related modulation of network pairs within the right hemisphere or between the right hemisphere and the rest of the brain, whereas others showed leftward segregation, reflecting preferential influence on within-left-hemisphere interactions.

**Figure 3.**
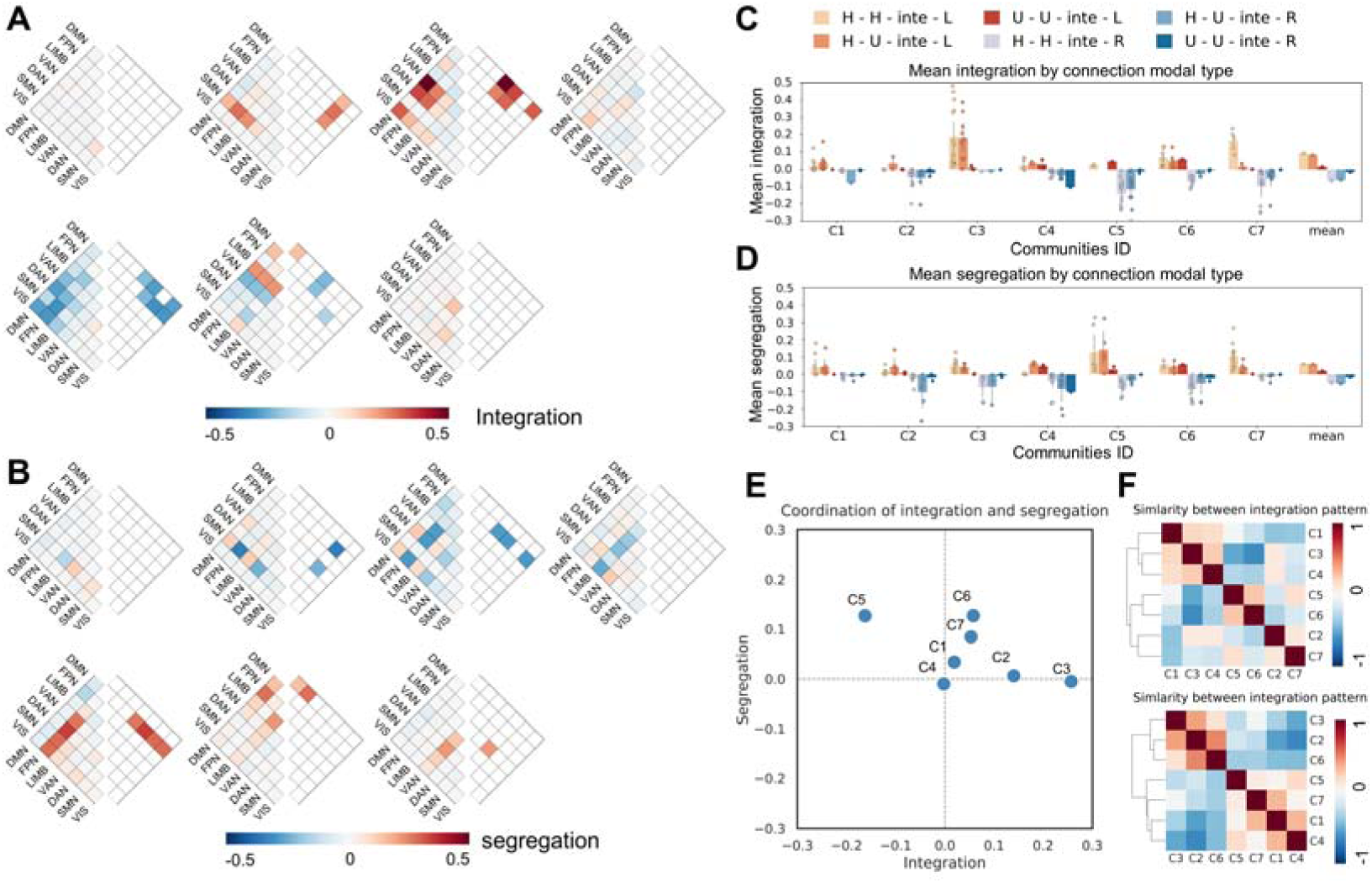
Arousal-modulated communities architecture exhibit community-specific hemispheric asymmetry. **(A)** Integration indices for each network pair revealed significant leftward or rightward deviations from a spatial permutation null, indicating that arousal differentially modulates between- and within-hemisphere interactions for specific network pairs. **(B)** Segregation indices identified network pairs showing hemisphere-specific strengthening of within-hemisphere connectivity, further demonstrating that lateralization is localized rather than global. **(C–D)** Community-averaged integration and segregation values showed that hemispheric biases vary across communities and are not determined solely by connection modal type, underscoring the community-specific nature of the asymmetry. **(E)** Aggregating all significantly lateralized network pairs, each community exhibited a distinct integration–segregation profile, revealing unique hemispheric signatures across communities. **(F)** Low similarity among communities’ integration and segregation patterns confirmed that arousal imposes multiple, community-specific forms of hemispheric asymmetry, rather than a single unified left- or right-dominant pattern.

Averaging indices across connection modal types reinforced this community-specific patterning (Fig. 3C–D). Notably, identical connection modal types (e.g., H-H) could show leftward bias in one community but rightward bias in another, demonstrating that lateralization is tied to the community architecture rather than to the canonical network pair class. Aggregating all significantly lateralized network pairs, each community expressed a unique lateralization signature (Fig. 3E), and there was no clear cluster of lateralization patterns among communities (Fig. 3F), indicating heterogeneous, rather than unified, hemispheric influences of arousal on FC. Instead, arousal imprints distinct left–right biases across different communities.

The robustness of these hemispheric lateralization of community architecture was further supported by the aforementioned resampling validations (Fig. S2). For the mean integration and segregation indices across network pairs, the empirical values consistently fell within the center of the resampling distributions. Specifically, for the network-pair specific lateralization signatures, the directional biases observed in each community remained stable during resampling, with empirical values largely aligned with the center of their respective distributions. This high degree of consistency confirms that the reported community-specific asymmetry is a stable and representative feature of arousal-modulated organization, ensuring that our findings are not skewed by specific sample compositions or outliers.

### Gradients of community affiliation entropy and the centripetal lateralization architecture of community affiliation

Having established that arousal-modulated functional connections can be organized into modular communities with distinct hemispheric lateralization at the network level, we next explored the nodal-level mapping of these communities and the substantial variability in how regions participate in arousal-modulated communities. For each region, we computed its community affiliation for each brain region, defined as the proportion of edges connected to that region that were assigned to each of the seven communities, separately for LL, LR, RL, and RR edges (Fig. 4 A–B).

**Figure 4.**
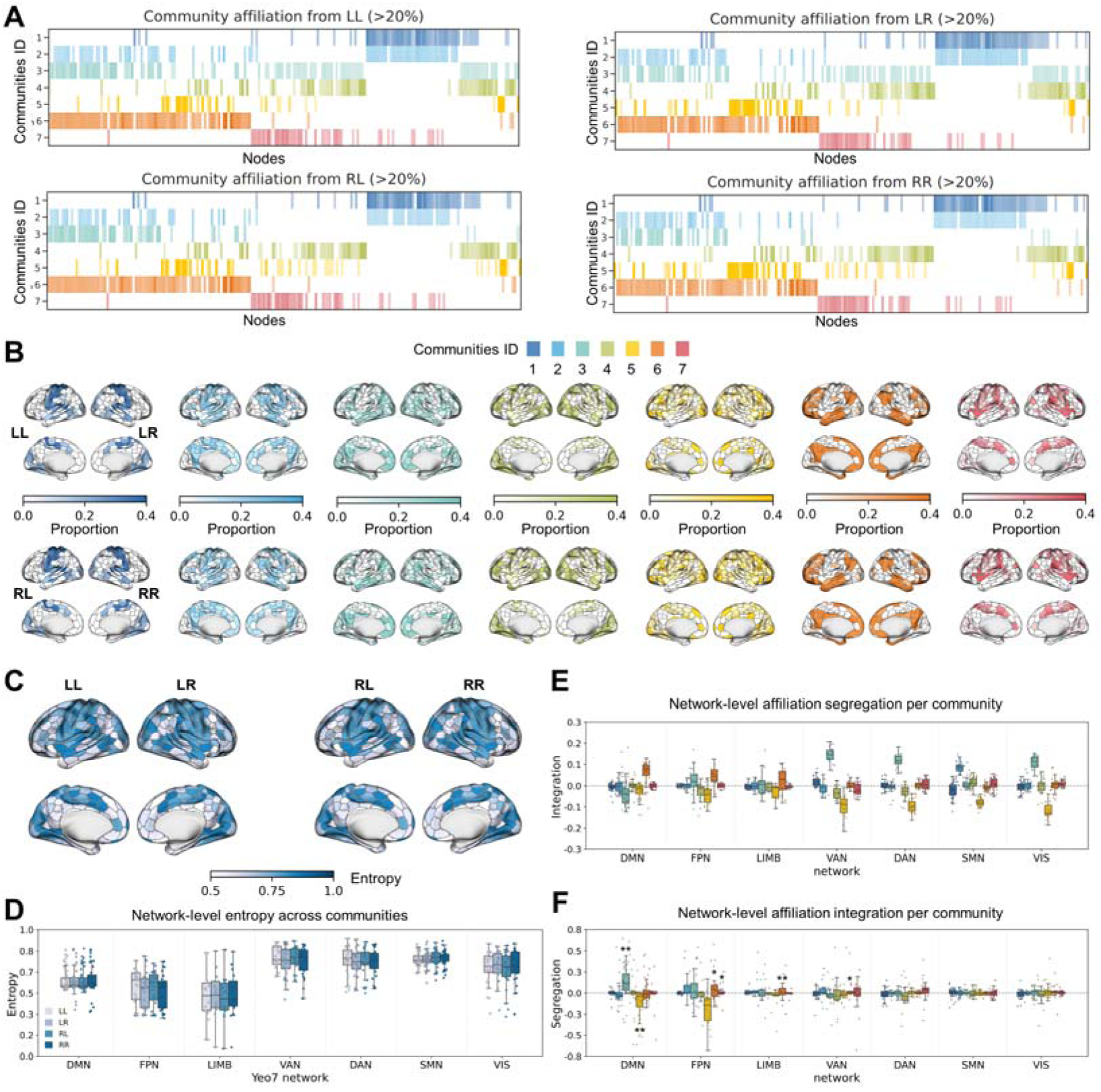
Characterizing the spatial distribution, entropy, and hemispheric divergence of regional community affiliation. **(A–B)** Nodal-level community affiliation matrices, computed separately for LL, LR, RL, and RR edges, showed substantial heterogeneity in how nodes distribute their arousal–tvFC coupling across the seven communities, with distinct patterns emerging across canonical networks. For visualization purposes, Figure A is restricted to displaying values where the proportion exceeds 0.2. **(C–D)** Region-level community affiliation entropy revealed a systematic network gradient, in which heteromodal systems displayed more selective participation, whereas unimodal network showed broader, more distributed engagement across communities (t(798) = -23.81, p = 4.24×10□^95^) **(E)** No integration bias was detected in any community. **(F)** Significant leftward segregation biases were identified within specific communities (communities 3, 5, 6, and 7). These asymmetries were primarily localized in regions belonging to the DMN, FPN, LIMB, and VAN. The color of each box corresponds to the community identity. Statistical significance: * p_FDR < 0.01; ** p_FDR < 0.001.

Next, we investigated the nodal-level community participation flexibility by calculating the community affiliation entropy for each region. Nodal-level affiliation entropy showed a clear network gradient (Fig. 4C–D). Default modal network (DMN), frontoparietal network (FPN), and limbic (LIMB) regions exhibited lower entropy, indicating focused participation in fewer arousal-modulated communities. In contrast, dorsal attention network (DAN), somatomotor network (SMN), and visual (VIS) regions showed broader participation across communities (t(798) = -23.81, p = 4.24×10 □_95_).

To evaluate lateralization at the nodal level, we derived metrics of segregation and integration based on these affiliation profiles. We observed a common organizational principle across several key communities: while no significant hemispheric bias in integration was detected in any community (Fig. 4E), Communities 3, 5, 6, and 7 exhibited robust leftward segregation biases (Fig. 4F). This dissociation between segregation and integration reveals a pronounced centripetal lateralization architecture. A detailed examination of the regional affiliation profiles indicated a consistent directional bias in signal flow across these communities: whereas left-hemisphere projections were predominantly intra-hemispheric (LL > LR), right-hemisphere inputs were biased contralaterally toward the left side (RL > RR). These findings suggest that these communities collectively represent a “left-hemisphere centripetal architecture”, where the left hemisphere serves as a preferential convergence of arousal-modulated signals, preferentially aggregating both ipsilateral and contralateral inputs.

The robustness of these observations was further supported by the aforementioned resampling validations (Fig. S3). Regarding the hemispheric lateralization of affiliation profiles, the directional biases observed across regions and communities remained highly stable during resampling, with empirical values consistently aligned with the center of their respective distributions. This high degree of consistency confirms that the reported centripetal affiliation architecture is a stable and representative feature of arousal-modulated lateralization, ensuring that our findings are not skewed by specific sample compositions or outliers.

### Arousal-tvFC coupling lateralization arises from spatial heterogeneity rather than mean shifts

In the preceding analyses, we mainly focused on the lateralization of the organizational patterns of the decomposed communities at the network-pairs and nodal levels. However, how the strength of the arousal–tvFC coupling is spatially distributed and whether lateralization exists in this distribution remains elusive. We therefore next examined the spatial distribution of arousal–tvFC coupling strength and its lateralization properties.

To test whether hemispheric biases reflected simple shifts in mean arousal–tvFC coupling strength, we compared mean integration and segregation across communities and whole connectome. Although the inter-community difference for segregation was statistically significant, the mean difference between communities was very small. This indicates limited hemispheric imbalance.

Although the mean arousal modulation on FC showed no significant lateralization, prior results suggested that its spatial pattern is highly community specific. We therefore hypothesized that the key information lies in the spatial heterogeneity (distribution gradient) of the modulation strength, not the overall strength. We therefore quantified spatial heterogeneity by ranking network pairs along a “lateralization axis” (Yang et al., 2025) and computing the slope of integration or segregation values along this axis. All communities showed slopes significantly steeper than the whole-brain baseline (all p_FDR < 0.001; Fig. 5B), demonstrating that lateralization arises from spatial heterogeneity—not uniform shifts.

**Figure 5.**
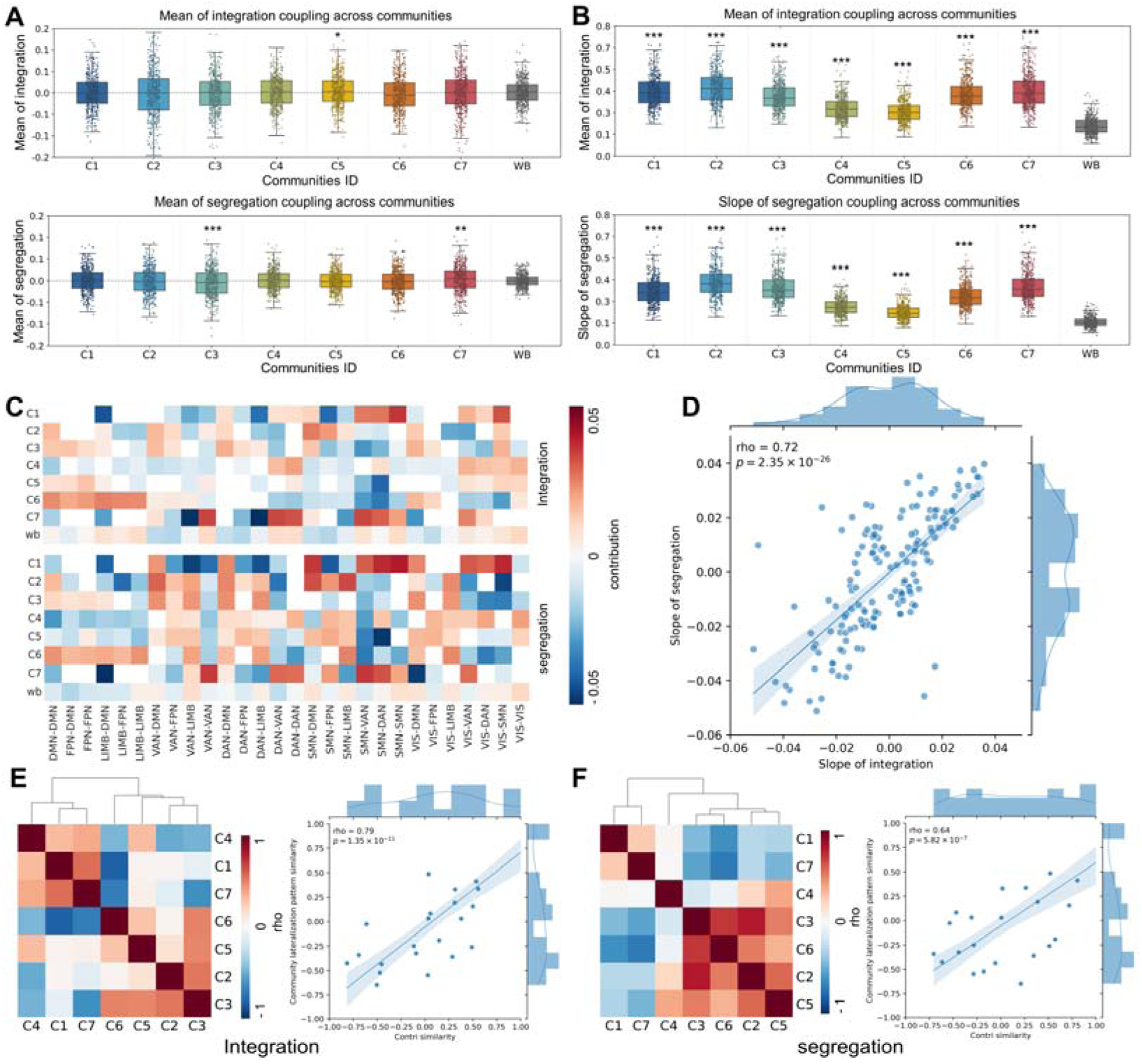
Spatial heterogeneity—not mean shifts—drives arousal-modulated hemispheric asymmetry. **(A)** Mean integration and segregation values showed no consistent hemispheric bias across individual communities or at the whole-brain (WB) level, indicating minimal imbalance in overall modulation strength. **(B)** Spatial heterogeneity, quantified as the slope of integration or segregation values ranked along a lateralization axis, was significantly steeper in every community compared with the whole-brain baseline (all p_FDR < 0.0001). These effects demonstrate that hemispheric asymmetry arises from spatially patterned variation, not from uniform shifts in mean modulation. **(C)** Leave-one-out analyses revealed that this heterogeneity reflects distributed contributions from many network pairs, rather than being driven by a few extreme edges. **(D)** Contribution patterns for integration and segregation were strongly correlated (rho ≈ 0.72, p = 2.35×10^-26^), indicating coordinated spatial organization across metrics. **(E–F)** Similarity in contribution patterns between communities was positively associated with similarity in communities’ intrinsic structure (integration: rho ≈ 0.79, p = 1.35×10^-11^; segregation: rho ≈ 0.64, p = 5.82×10^-7^), showing that spatial heterogeneity is constrained by each community’s underlying connectivity architecture. Statistical note: * p_FDR < 0.01; ** p_FDR < 0.001; *** p_FDR < 0.0001.

Leave-one-out analyses showed that heterogeneity does not depend on a small set of extreme network pairs. Instead, most pairs contributed modestly, producing broad, community-specific gradients (Fig. 5C). Contribution patterns for integration and segregation were highly correlated (rho ≈ 0.72, p = 2.35×10^-26^). Furthermore, similarity in contribution patterns tracked similarity in intrinsic community structure (integration: rho ≈ 0.79, p = 1.35×10^-11^; segregation: rho ≈ 0.64, p = 5.82×10^-7^), indicating that spatial heterogeneity is shaped by underlying connectivity architecture (Fig. 5E–F).

Thus, arousal-driven lateralization is best understood as structured spatial heterogeneity within communities, rather than gross hemispheric dominance.

The robustness of these observations was further supported by the aforementioned resampling validations (Fig. S4). For the mean integration and segregation indices, the empirical values consistently fell within the center of the resampling distributions, reinforcing the absence of a uniform hemispheric mean shift across the population. Conversely, for the spatial heterogeneity metrics, the slopes observed in each community remained stable during resampling, with the empirical values largely aligned with the center of their respective distributions. This high degree of consistency confirms that the reported spatial heterogeneity is a stable and representative feature of arousal-modulated lateralization, ensuring that our findings are not skewed by specific sample compositions or outliers.

### Community structure and hemispheric asymmetry are preserved during movie watching

To assess whether these organizational principles generalize beyond the resting state, we applied the same analytic pipeline to naturalistic movie-watching data from the same participants.

The seven-community structure was highly preserved across paradigms. After aligning communities using Hungarian algorithm (Kuhn, 1955), we found that the overall community structure showed consistent correspondence (average dice ≈ 0.46; Fig. 6A–B). One heteromodal-dominated community (community 6) showed near-perfect correspondence (rho ≈ 0.94), suggesting that its arousal sensitivity is largely independent of external stimulation. Other communities also demonstrated moderate cross-paradigm similarity, with only two showing low similarity—likely reflecting context-dependent modulation under rich sensory input.

**Figure 6.**
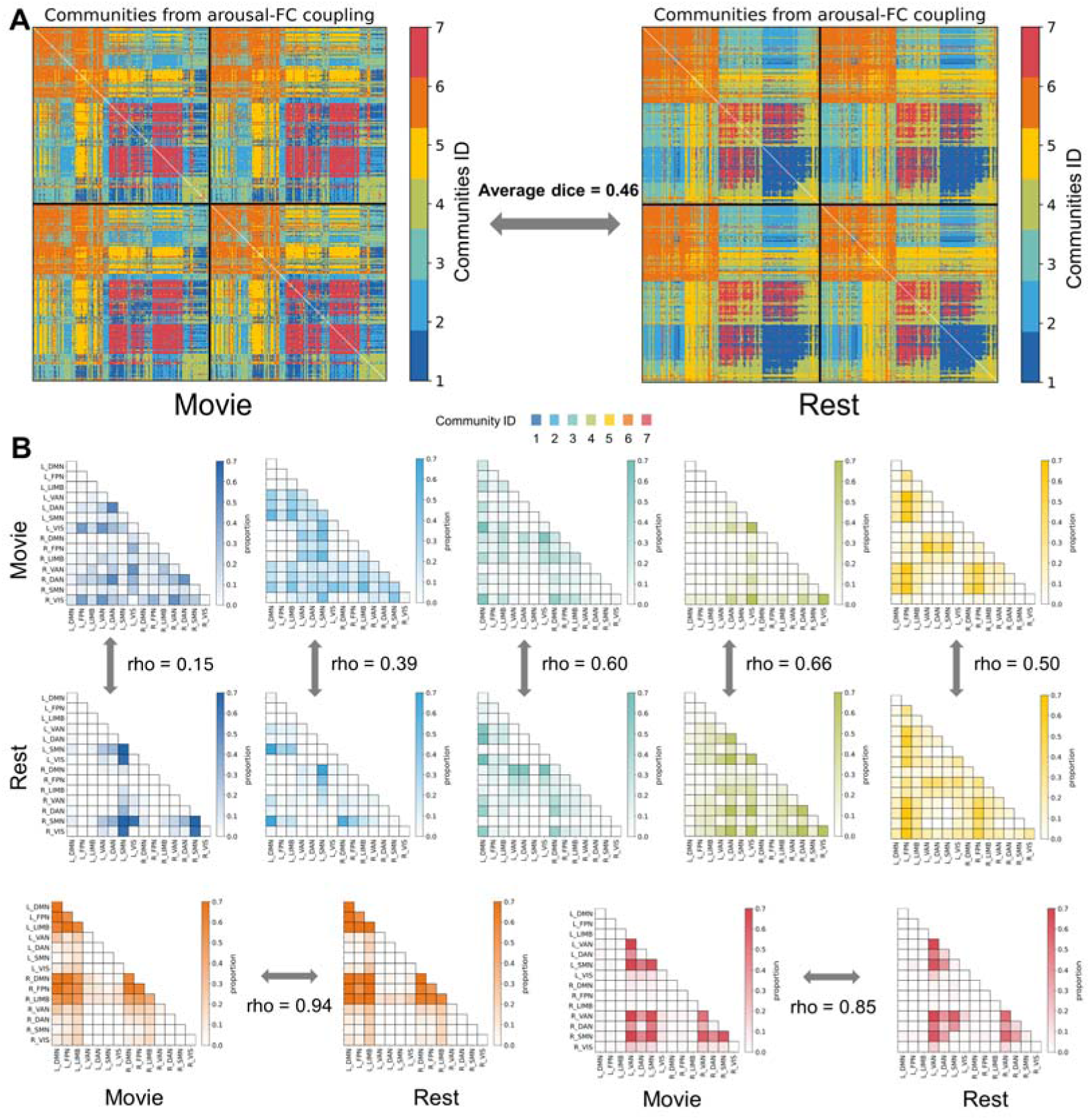
Community structure and hemispheric asymmetry of arousal–tvFC coupling are preserved across resting state and movie watching. **(A)** Communities derived from rest and movie data were aligned using Hungarian algorithm, revealing robust correspondence between paradigms (average dice ≈ 0.46). **(B)** Network-pair composition profiles for each community were strongly correlated across paradigms (mean rho ≈ 0.58), indicating that the modular organization of arousal–tvFC coupling is stable across cognitive contexts.

These findings indicate that the modular, asymmetric organization of arousal–tvFC coupling is not specific to rest but reflects intrinsic principles that persist across cognitive contexts.

## Discussion

In this study, we investigated how moment-to-moment fluctuations in arousal shape large-scale functional connectivity in the awake human brain. By combining high-field fMRI with concurrent pupillometry, we quantified arousal–tvFC coupling at the level of individual edges and showed that arousal does not exert a uniform or diffuse influence across the connectome. Instead, it modulates connectivity through a set of seven distinct communities, each defined by characteristic network compositions and hemispheric patterns. These findings demonstrate that fluctuations in arousal, even within stable wakefulness, impose a structured and asymmetric organization on whole-brain functional interactions.

Although arousal is often conceptualized as a global modulatory state, our results show that its impact on tvFC is highly organized. Specifically, we show the presence of low-dimensional, reproducible communities suggests that arousal modulates the connectome through structured spatial patterns rather than homogeneous gain modulation. We hypothesize that this structured macroscopic architecture reflects the differentiated projection patterns of subcortical neuromodulatory systems, such as the locus coeruleus–noradrenergic pathway (Aston-Jones & Cohen, 2005; Jordan, 2024) and thalamus (Magnin et al., 2010; Lewis et al., 2015; Liu et al., 2018). This organized pattern of modulation further supports the view advocated in recent years that arousal states should be viewed as possessing spatiotemporal dynamics and regional complexity (Siclari & Tononi, 2017; Nir & De Lecea, 2023), and is strongly supported by prior work showing that spontaneous arousal fluctuations influence distributed cortical responses in selective ways (Reimer et al., 2014; McGinley et al., 2015). The hierarchical community pattern—where unimodal interactions cluster into fewer patterns while heteromodal systems participate broadly—is compatible with theories that place heteromodal cortex at the apex of large-scale integrative gradients (Mesulam, 1998; Margulies et al., 2016). Together, these observations suggest that the arousal-sensitive connectome reflects not merely regional susceptibility but an intrinsic network-level architecture, in which state-dependent modulation is aligned with established large-scale organizational principles.

A major goal of this work was to determine whether arousal imposes systematic hemispheric asymmetry on FC. We found clear evidence that it does, but in a community-specific rather than global manner. Certain communities showed rightward integration, others leftward segregation, and others no significant bias—indicating that hemispheric asymmetry emerges from the organization of arousal-sensitive connectivity motifs rather than a single overarching hemispheric dominance. This distributed asymmetry resonates with converging evidence across species: unihemispheric vigilance in birds and marine mammals (Rattenborg et al., 2000; Lyamin et al., 2016; Mascetti, Gian Gastone, 2016; Reicher et al., 2021; Fenk et al., 2023; Libourel et al., 2023), asymmetric EEG signatures related to vigilance in humans (Tamaki et al., 2016), and classic findings of right-lateralized alerting and reorienting functions (Heilman & Abell, 1980; Sturm & Willmes, 2001; Shulman et al., 2010; Corbetta & Shulman, 2011). This suggests that hemispheric specialization may arise from distinct modes of arousal-modulated reconfiguration rather than fixed structural asymmetries.

Despite the robustness of these hemispheric differences, mean integration and segregation of arousal-tvFC coupling strength across the entire community or whole brain showed minimal global lateralization. Instead, arousal-driven asymmetry manifested as spatially heterogeneous gradients within communities. Arousal does not uniformly shift connectivity toward one hemisphere; rather, it selectively amplifies lateralization in specific connectivity motifs. This distributed, gradient-like pattern complements recent work highlighting macroscale cortical gradients and manifold structure as fundamental organizational principles (Margulies et al., 2016; Huntenburg et al., 2018). We further found that communities with similar intrinsic topology exhibited similar heterogeneity signatures, suggesting that baseline connectivity architecture constrains how arousal shapes hemispheric interactions. This provides a mechanistic explanation for why traditional hemisphere-level metrics often obscure arousal-modulated asymmetries—these effects are expressed not as global biases but as structured, topology-dependent gradients.

Another important observation is the stability of arousal–tvFC organization across cognitive contexts. While the overall spatial layout of the seven-community architecture showed moderate reorganization between resting-state and naturalistic movie-watching, specific motifs—most notably communities 6 and 7—demonstrated near-perfect correspondence across paradigms. This robustness aligns with prior evidence that intrinsic connectivity organization persists across tasks (Cole et al., 2014; Krienen et al., 2014) and that spontaneous fluctuations in arousal modulate cortical dynamics even during rich sensory stimulation (Tanner et al., 2023). By demonstrating that arousal-driven network modulation generalizes across both internally and externally oriented states, our findings indicate that arousal acts as a stable organizing axis of large-scale brain communication, rather than merely a background physiological fluctuation.

Despite the important findings of this study, several limitations should be noted. First, to ensure a mathematically rigorous assessment of hemispheric asymmetry, our analysis was restricted to a symmetric cortical parcellation. Consequently, while we demonstrate that arousal-modulated connectivity follows a structured macroscopic architecture, we did not explicitly analyze the subcortical nuclei hypothesized to drive these patterns. We hypothesize that the presence of these low-dimensional cortical communities reflects coordinated motifs rather than a homogeneous gain modulation, potentially mirroring the differentiated projection patterns of subcortical neuromodulatory systems. For instance, the locus coeruleus–noradrenergic pathway (Chandler et al., 2014; Schwarz & Luo, 2015) and thalamus (Hwang et al., 2017; Shine, 2019; Müller et al., 2020; Shine et al., 2023) possess extensive yet non-uniform projections that may anchor the community-specific and hemispherically asymmetric patterns observed here. In the absence of direct subcortical-cortical integration in our current framework, this link remains hypothetical. Future investigations incorporating high-resolution subcortical data will be essential to empirically bridge the gap between these large-scale cortical communities and the underlying physiological scaffolding of the Ascending Reticular Activating System (ARAS). Second, while pupillometry is a well-established and accessible proxy for central arousal (Joshi & Gold, 2020), it remains an indirect peripheral marker. Integrating these findings with concurrent EEG will be essential to provide a more granular, multi-modal characterization of arousal states. Third, we utilized sliding-window FC to track time-varying connectivity. Although this is a robust and widely validated technique, employing alternative dynamic approaches with higher temporal resolution—such as edge FC (Faskowitz et al., 2020), multiplication of temporal derivatives (Shine et al., 2015) or dynamic conditional correlation (Lindquist et al., 2014)—may offer complementary perspectives on the transient dynamics of arousal-modulated states. Fourth, the generalizability of our approach to external cohorts warrants caution regarding pupillary data integrity. In contexts where high-fidelity eye-tracking is technically demanding—such as in clinical settings involving patients with restricted compliance or in naturalistic fMRI studies—the prevalence of blink artifacts and signal dropouts may bias the estimation of arousal-modulated states. Excessive reliance on data interpolation in such cases could artificially smooth temporal fluctuations, leading to an overestimation of community stability. Future applications should therefore prioritize high-frequency sampling and potentially incorporate multi-modal physiological features (e.g., respiratory or cardiac signals) to cross-validate arousal dynamics when pupillary data is suboptimal (Meissner et al., 2023; Bolt et al., 2025; Weijs et al., 2025). Fifth, the findings reported here were derived exclusively from ultra-high-field (7T) imaging data. The superior BOLD sensitivity of 7T fMRI was instrumental in resolving the fine-scale community architecture of arousal–tvFC coupling, which involves subtle signals that may be challenging to detect at lower field strengths. Given that 3T remains the most common parameter for neuroimaging research and clinical applications, future investigations are needed to determine the extent to which these organizational principles generalize to standard field strength data. Validating these communities in large-scale 3T datasets will be essential to establish their broader applicability across different imaging environments. Sixth, our findings were derived using a single high-resolution cortical parcellation. While the specific choice of atlas can influence fine-grained regional connectivity, it is important to note that our primary conclusions—such as hemispheric asymmetries and community-level preferences—were identified and interpreted at the macroscopic network and system level. By aggregating signals across broad functional systems, this approach likely mitigates the dependency on precise regional boundary definitions. Nevertheless, future studies employing alternative parcellation schemes would be valuable to further confirm that these organizational principles are not specific to the current atlas but represent a generalizable feature of the arousal-modulated connectome.

In summary, moment-to-moment fluctuations in arousal modulate functional connectivity through a small number of structured connectivity communities, each with distinct hemispheric characteristics. These asymmetries arise not from global shifts but from spatially heterogeneous gradients embedded within community structure. The reproducibility of these communities across resting state and naturalistic stimulation suggests that they reflect stable and intrinsic principles by which arousal dynamically shapes large-scale brain interactions during wakefulness.

## Methods and Materials

### Participants and datasets

We used data from the Human Connectome Project (HCP) 7T dataset, which includes resting-state fMRI and naturalistic movie-watching runs with simultaneous eye tracking (Van Essen et al., 2013). Across participants, up to four resting runs and four movie runs were available, each approximately 15–16 minutes in length. All procedures were approved by the Washington University Institutional Review Board, and written informed consent was obtained from all participants.

### MRI acquisition and preprocessing

All imaging was collected on a Siemens 7T scanner (TR = 1 s, TE = 22.2 ms, voxel size = 1.6 mm isotropic, multiband factor = 5, 900 volumes/run). We used the HCP minimally preprocessed data (Glasser et al., 2013), followed by additional denoising: linear detrending; regression of 24 head-motion parameters; regression of white matter and CSF signals; band-pass filtering (0.01–0.1 Hz); scrubbing of frames with FD > 0.2 mm; and interpolation across removed frames.

### ROI parcellation

Analyses were performed using a 400-ROI symmetric parcellation with explicitly matched left–right homologues (Yan et al., 2023). Vertexwise BOLD signals were averaged within each ROI to obtain regional time series. All subsequent hemispheric analyses relied on this explicit homotopic structure.

### Eye tracking preprocessing

Pupil diameter was extracted from the raw eye-tracking stream and cleaned following established procedures (Gonzalez-Castillo et al., 2022): Removal of samples outside MRI acquisition; detection of blinks and short missing segments (<1 s); linear interpolation across missing segments; removal of brief physiologically implausible excursions (<1 ms within long closures); smoothing with a 200 ms Hanning window; down-sampling to 1 Hz to match the temporal resolution of the fMRI data. The resulting time series served as a continuous arousal index for each run.

### Quality control

The final analyzed sample for the resting-state consisted of N = 139 healthy participants (mean age = 29.1±3.5 years, 77 female). Runs were excluded if (a) more than 20% of frames exceeded motion thresholds, (b) eye tracking did not cover the full fMRI time series, or (c) more than 90% of samples were classified as eye closure. After applying these criteria, 485 of the initial 723 scans were retained for analysis. The same quality-control pipeline was applied to the movie-watching dataset, yielding 513 usable scans out of the original 725. After rigorous quality control, 139 participants (485 runs) were retained for the final analysis. Detailed information on data retention and run distribution per participant is summarized in Figure S9.

### Time-varying functional connectivity

Time-varying FC between each pair of ROIs was estimated using sliding-window correlations: window length: 30 s; step size: 5 s. Within each window, Pearson correlation coefficients were computed and Fisher-z transformed. This procedure yielded a tvFC time series for each edge in each run.

### Arousal estimation from pupil size

Pupil data were processed using the same sliding-window parameters. The mean pupil size within each window was taken as an index of moment-to-moment arousal level.

### Estimating arousal–tvFC coupling

For each functional connection, arousal–tvFC coupling was defined as the Pearson correlation between its time-varying FC and the pupil-derived arousal fluctuations across windows. Thus, each run produced a 400 × 400 symmetric matrix of coupling values, later vectorized into edgewise features.

These matrices were concatenated across runs to form the dataset used for community detection and all subsequent analyses.

### Community detection on arousal–tvFC coupling

To identify brain regions sharing similar arousal-related modulation profiles, we performed community detection on the edgewise coupling values. Prior to clustering, coupling values were z-scored across runs to ensure comparability. In this analytical framework, brain edges were treated as observations, while the individual runs served as feature dimensions, effectively representing each edge by its unique across-run coupling motif.

We employed the k-means clustering algorithm (Euclidean distance) to explore a range of cluster solutions from K = 2 to 15. To ensure the stability of the results and avoid local optima, each K was repeated 250 times with random initializations. The optimal number of clusters was determined by evaluating clustering quality and reproducibility (e.g., maximizing silhouette stability). It is important to clarify that “communities” in this context refer to clusters of edges that exhibit similar arousal-modulation motifs within a high-dimensional feature space, rather than topological modules typically derived from graph-theoretic algorithms like modularity maximization (Blondel et al., 2008). This procedure consistently identified seven distinct communities, each representing an arousal-sensitive connectivity motif.

### Mapping communities to networks and computing entropy

Each edge was assigned to one of seven communities. Edges were then mapped to Yeo’s 7 canonical networks (DMN, FPN, LIMB, VAN, DAN, SMN and VIS) (Thomas Yeo et al., 2011).

For each network pair*(i,j)* we computed the proportion *p_i,j,k_* of its edges belonging to each community *k*. Shannon entropy *H_i,j_* quantified how broadly a network pair participated across communities, calculated as:

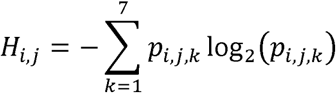

A higher *H_i,j_* indicates a broader, more uniform participation across the seven communities, while a lower *H_i,j_* indicates that the edges are primarily concentrated in a few communities.

Additionally, the dominant community for each network pair is defined as

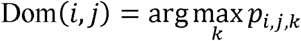

Entropy and dominant-community assignments jointly characterized both the diversity and primary affiliation of each network pair within the community structure.

Nodal-level community affiliation entropy A*_r_* quantified how broadly each region *r* participated across communities. The entropy was calculated using the proportion q *_r_*_,*k*_ of all incident edges of the region *r* belonging to the community *k*:

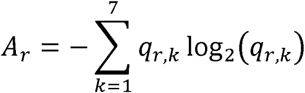

### Quantifying hemispheric asymmetry: integration and segregation indices

To evaluate hemispheric biases in arousal–tvFC coupling, we categorized all functional edges into four types based on their nodal locations: LL (within left hemisphere), RR (within right hemisphere), LR (left-to-right), and RL (right-to-left). Following established frameworks for lateralization (Gotts et al., 2013), we calculated two complementary indices to capture the nature of this asymmetry.

The integration index provides a measure of the overall hemispheric dominance of arousal-modulated connections. A positive value indicates that arousal-modulated edges are preferentially concentrated in the left hemisphere (encompassing both its intra-hemispheric and commissural connections) relative to the right. It is defined as:

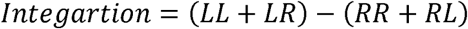

The segregation index assesses whether arousal preferentially modulates local, intra-hemispheric communication versus long-range, inter-hemispheric communication. A positive value reflects a “segregated” left-hemisphere bias, where arousal strengthens connections within the left hemisphere more than it strengthens its communication with the contralateral hemisphere. It is defined as:

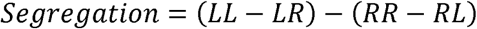

Indices were computed for each network pair, each community (weighted by edge count), and all significantly lateralized subsets. Statistical significance was assessed relative to a null model (see below).

### Spatial heterogeneity of lateralization

To quantify the spatial distribution characteristics of the arousal-tvFC coupling strength features (e.g., integration, segregation) within each community *k*, we first projected the edgewise coupling matrix for each participant onto the network-pair level, following the same procedure described in the previous section. The average coupling value of each network pair (*i,j*) within each community *k* is C*_i,j,k_* Network pairs were then sorted by their lateralization values to define the “lateralization axis” (Yang et al., 2025).

Spatial heterogeneity was quantified by performing a linear regression of the network-pair coupling values against their rank along the axis:

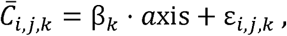

the regression slope *β_k_* indexed the spatial heterogeneity, where a larger | *β_k_*| indicates a greater difference in lateralization value within the community.

The influence of each individual network pair (*i,j*) on the overall spatial heterogeneity β*_k_* was assessed using a leave-one-out method. The contribution D*_i,j,k_* was defined as the difference between the new slope *β̄_i,ijk_* (after removing the pair) and the original slope β*_k_*:

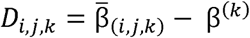

The sign of *D_i,j,k_* indicates its modulatory role: positive values enhance spatial heterogeneity, whereas negative values reduce it.

### Null model for hemispheric asymmetry

To determine whether observed lateralization exceeded chance, we construct a null model for hemispheric asymmetry.

For each run, ROI indices were permuted identically for rows and columns, preserving matrix symmetry and degree distribution while disrupting hemispheric structure. 10,000 permutations were performed. For each iteration, clustering and asymmetry indices were recomputed. p-values were FDR-corrected across comparisons.

### Cross-paradigm validation using movie watching

To assess the context-independence of arousal–tvFC organization, we applied all analyses to movie-watching runs. Community structures from rest and movie data were matched using Hungarian assignment (Kuhn, 1955). The overall similarity between the community architectures derived from the two paradigms was quantified by the mean Dice coefficient. Community-level correspondence was quantified by Spearman correlation between network-pair community profiles across paradigms.

### Robustness and validation

To ensure the stability and generalizability of our findings, we performed extensive robustness analyses across multiple biological scales. We first employed two complementary resampling strategies—500-iteration split-half reliability tests and participant-level resampling—with the entire analytical pipeline re-executed for each iteration. The stability of community partitions was quantified using Dice coefficients, while the reliability of hemispheric asymmetry indices was assessed by their average deviation from the full-dataset estimates. Crucially, we further confirmed that these organizational patterns were not driven by non-neural confounds, as the identified community architecture and lateralization remained highly stable even after explicitly controlling for head motion and the global signal in the arousal–tvFC coupling model. A series of sensitivity analyses regarding sliding-window parameters, temporal lags, and alternative pupillometry preprocessing pipelines further supported the robustness of our results. Detailed procedures and supporting results for these validations are provided in Supplementary Methods S1–S6 and Figures S1–S9.

## Data availability

Raw and preprocessed HCP data can be accessed at https://db.humanconnectome.org/. Source data to replicate the results of the study are openly available at https://github.com/kongxy6478/Arousal-modulates-functional-connectivity.

## Code availability

All analyses were implemented in Python (NumPy, SciPy, scikit-learn) using custom scripts. Visualization was performed with Matplotlib, Seaborn, and Surfplot. Computer codes used to calculate the communities, analyse results and reproduce the figures of the study are openly available at https://github.com/kongxy6478/Arousal-modulates-functional-connectivity.

## Acknowledgment

This work is supported by the National Natural Science Foundation of China (Nos. T2325006, 82021004), STI 2030-Major Projects (Nos.2021ZD0201701, 2021ZD0200500); the Fundamental Research Funds for the Central Universities (No. 2233200020).

## Reference

Aston-Jones, G., & Cohen, J. D. (2005). AN INTEGRATIVE THEORY OF LOCUS COERULEUS-NOREPINEPHRINE FUNCTION: Adaptive Gain and Optimal Performance. In Annual Review of Neuroscience (Vol. 28, Issue Volume 28, 2005, pp. 403–450). Annual Reviews. 10.1146/annurev.neuro.28.061604.135709

Banks, M. I., Krause, B. M., Endemann, C. M., Campbell, D. I., Kovach, C. K., Dyken, M. E., Kawasaki, H., & Nourski, K. V. (2020). Cortical functional connectivity indexes arousal state during sleep and anesthesia. NeuroImage, 211, 116627. 10.1016/j.neuroimage.2020.116627

Blondel, V. D., Guillaume, J.-L., Lambiotte, R., & Lefebvre, E. (2008). Fast unfolding of communities in large networks. Journal of Statistical Mechanics: Theory and Experiment, 2008(10), P10008. 10.1088/1742-5468/2008/10/P10008

Bolt, T., Wang, S., Nomi, J. S., Setton, R., Gold, B. P., deB. Frederick, B., Yeo, B. T. T., Chen, J. J., Picchioni, D., Duyn, J. H., Spreng, R. N., Keilholz, S. D., Uddin, L. Q., & Chang, C. (2025). Autonomic physiological coupling of the global fMRI signal. Nature Neuroscience, 28(6), 1327–1335. 10.1038/s41593-025-01945-y

Breeden, A. L., Siegle, G. J., Norr, M. E., Gordon, E. M., & Vaidya, C. J. (2017). Coupling between spontaneous pupillary fluctuations and brain activity relates to inattentiveness. European Journal of Neuroscience, 45(2), 260–266. 10.1111/ejn.13424

Chandler, D. J., Gao, W.-J., & Waterhouse, B. D. (2014). Heterogeneous organization of the locus coeruleus projections to prefrontal and motor cortices. Proceedings of the National Academy of Sciences, 111(18), 6816–6821. 10.1073/pnas.1320827111

Chow, H. M., Horovitz, S. G., Carr, W. S., Picchioni, D., Coddington, N., Fukunaga, M., Xu, Y., Balkin, T. J., Duyn, J. H., & Braun, A. R. (2013). Rhythmic alternating patterns of brain activity distinguish rapid eye movement sleep from other states of consciousness. Proceedings of the National Academy of Sciences, 110(25), 10300–10305. 10.1073/pnas.1217691110

Cole, M. W., Bassett, D. S., Power, J. D., Braver, T. S., & Petersen, S. E. (2014). Intrinsic and Task-Evoked Network Architectures of the Human Brain. Neuron, 83(1), 238–251. 10.1016/j.neuron.2014.05.014

Corbetta, M., & Shulman, G. L. (2011). Spatial Neglect and Attention Networks. In Annual Review of Neuroscience (Vol. 34, Issue Volume 34, 2011, pp. 569–599). Annual Reviews. 10.1146/annurev-neuro-061010-113731

Damaraju, E., Tagliazucchi, E., Laufs, H., & Calhoun, V. D. (2020). Connectivity dynamics from wakefulness to sleep. NeuroImage, 220, 117047. 10.1016/j.neuroimage.2020.117047

Demertzi, A., Tagliazucchi, E., Dehaene, S., Deco, G., Barttfeld, P., Raimondo, F., Martial, C., Fernández-Espejo, D., Rohaut, B., Voss, H. U., Schiff, N. D., Owen, A. M., Laureys, S., Naccache, L., & Sitt, J. D. (2019). Human consciousness is supported by dynamic complex patterns of brain signal coordination. Science Advances, 5(2), eaat7603. 10.1126/sciadv.aat7603

Faskowitz, J., Esfahlani, F. Z., Jo, Y., Sporns, O., & Betzel, R. F. (2020). Edge-centric functional network representations of human cerebral cortex reveal overlapping system-level architecture. Nature Neuroscience, 23(12), 1644–1654. 10.1038/s41593-020-00719-y

Fenk, L. A., Riquelme, J. L., & Laurent, G. (2023). Interhemispheric competition during sleep. Nature, 616(7956), 312–318. 10.1038/s41586-023-05827-w

Glasser, M. F., Sotiropoulos, S. N., Wilson, J. A., Coalson, T. S., Fischl, B., Andersson, J. L., Xu, J., Jbabdi, S., Webster, M., Polimeni, J. R., Van Essen, D. C., & Jenkinson, M. (2013). The minimal preprocessing pipelines for the Human Connectome Project. NeuroImage, 80, 105–124. 10.1016/j.neuroimage.2013.04.127

Gonzalez-Castillo, J., Fernandez, I. S., Handwerker, D. A., & Bandettini, P. A. (2022). Ultra-slow fMRI fluctuations in the fourth ventricle as a marker of drowsiness. NeuroImage, 259, 119424. 10.1016/j.neuroimage.2022.119424

Gotts, S. J., Jo, H. J., Wallace, G. L., Saad, Z. S., Cox, R. W., & Martin, A. (2013). Two distinct forms of functional lateralization in the human brain. Proceedings of the National Academy of Sciences, 110(36), E3435–E3444. 10.1073/pnas.1302581110

Heilman, K. M., & Abell, T. V. D. (1980). Right hemisphere dominance for attention: The mechanism underlying hemispheric asymmetries of inattention (neglect). Neurology, 30(3), 327–327. 10.1212/WNL.30.3.327

Huang, Z., Zhang, J., Wu, J., Mashour, G. A., & Hudetz, A. G. (2020). Temporal circuit of macroscale dynamic brain activity supports human consciousness. Science Advances, 6(11), eaaz0087. 10.1126/sciadv.aaz0087

Huntenburg, J. M., Bazin, P. L., & Margulies, D. S. (2018). Large-Scale Gradients in Human Cortical Organization. Trends Cogn Sci, 22(1), 21–31. https://doi.org/10/gcr9h4

Hwang, K., Bertolero, M. A., Liu, W. B., & D’Esposito, M. (2017). The Human Thalamus Is an Integrative Hub for Functional Brain Networks. The Journal of Neuroscience, 37(23), 5594–5607. 10.1523/JNEUROSCI.0067-17.2017

Jang, H., Mashour, G. A., Hudetz, A. G., & Huang, Z. (2024). Measuring the dynamic balance of integration and segregation underlying consciousness, anesthesia, and sleep in humans. Nature Communications, 15(1), 9164. 10.1038/s41467-024-53299-x

Jordan, R. (2024). The locus coeruleus as a global model failure system. Trends in Neurosciences, 47(2), 92–105. 10.1016/j.tins.2023.11.006

Joshi, S., & Gold, J. I. (2020). Pupil Size as a Window on Neural Substrates of Cognition. Trends in Cognitive Sciences, 24(6), 466–480. 10.1016/j.tics.2020.03.005

Krienen, F. M., Yeo, B. T. T., & Buckner, R. L. (2014). Reconfigurable task-dependent functional coupling modes cluster around a core functional architecture. Philosophical Transactions of the Royal Society B: Biological Sciences, 369(1653), 20130526. 10.1098/rstb.2013.0526

Kuhn, H. W. (1955). The Hungarian method for the assignment problem. Naval Research Logistics Quarterly, 2(1–2), 83–97. 10.1002/nav.3800020109

Lewis, L. D., Voigts, J., Flores, F. J., Schmitt, L. I., Wilson, M. A., Halassa, M. M., & Brown, E. N. (2015). Thalamic reticular nucleus induces fast and local modulation of arousal state. eLife, 4, e08760. 10.7554/eLife.08760

Libourel, P.-A., Lee, W. Y., Achin, I., Chung, H., Kim, J., Massot, B., & Rattenborg, N. C. (2023). Nesting chinstrap penguins accrue large quantities of sleep through seconds-long microsleeps. Science, 382(6674), 1026–1031. 10.1126/science.adh0771

Lindquist, M. A., Xu, Y., Nebel, M. B., & Caffo, B. S. (2014). Evaluating dynamic bivariate correlations in resting-state fMRI: A comparison study and a new approach. NeuroImage, 101, 531–546. 10.1016/j.neuroimage.2014.06.052

Liu, X., De Zwart, J. A., Schölvinck, M. L., Chang, C., Ye, F. Q., Leopold, D. A., & Duyn, J. H. (2018). Subcortical evidence for a contribution of arousal to fMRI studies of brain activity. Nature Communications, 9(1), 395. 10.1038/s41467-017-02815-3

Lloyd, B., De Voogd, L. D., Mäki-Marttunen, V., & Nieuwenhuis, S. (2023). Pupil size reflects activation of subcortical ascending arousal system nuclei during rest. eLife, 12, e84822. 10.7554/eLife.84822

Lyamin, O. I., Lapierre, J. L., Kosenko, P. O., Kodama, T., Bhagwandin, A., Korneva, S. M., Peever, J. H., Mukhametov, L. M., & Siegel, J. M. (2016). Monoamine Release during Unihemispheric Sleep and Unihemispheric Waking in the Fur Seal. Sleep, 39(3), 625–636. 10.5665/sleep.5540

Magnin, M., Rey, M., Bastuji, H., Guillemant, P., Mauguière, F., & Garcia-Larrea, L. (2010). Thalamic deactivation at sleep onset precedes that of the cerebral cortex in humans. Proceedings of the National Academy of Sciences, 107(8), 3829–3833. 10.1073/pnas.0909710107

Margulies, D. S., Ghosh, S. S., Goulas, A., Falkiewicz, M., Huntenburg, J. M., Langs, G., Bezgin, G., Eickhoff, S. B., Castellanos, F. X., Petrides, M., Jefferies, E., & Smallwood, J. (2016). Situating the default-mode network along a principal gradient of macroscale cortical organization. Proc Natl Acad Sci U S A, 20191114, 113(44), 12574–12579. https://doi.org/10/f89vsk

Mascetti, Gian Gastone, G. G. (2016). Unihemispheric sleep and asymmetrical sleep: Behavioral, neurophysiological, and functional perspectives. *Nature and Science of Sleep*, Volume 8, 221–238. 10.2147/NSS.S71970

McGinley, M. J., Vinck, M., Reimer, J., Batista-Brito, R., Zagha, E., Cadwell, C. R., Tolias, A. S., Cardin, J. A., & McCormick, D. A. (2015). Waking State: Rapid Variations Modulate Neural and Behavioral Responses. Neuron, 87(6), 1143–1161. 10.1016/j.neuron.2015.09.012

Meissner, S. N., Bächinger, M., Kikkert, S., Imhof, J., Missura, S., Carro Dominguez, M., & Wenderoth, N. (2023). Self-regulating arousal via pupil-based biofeedback. Nature Human Behaviour, 8(1), 43–62. 10.1038/s41562-023-01729-z

Mesulam, M. (1998). From sensation to cognition. Brain, 121(6), 1013–1052. 10.1093/brain/121.6.1013

Müller, E. J., Munn, B., Hearne, L. J., Smith, J. B., Fulcher, B., Arnatkevičiūtė, A., Lurie, D. J., Cocchi, L., & Shine, J. M. (2020). Core and matrix thalamic sub-populations relate to spatio-temporal cortical connectivity gradients. NeuroImage, 222, 117224. 10.1016/j.neuroimage.2020.117224

Nir, Y., & De Lecea, L. (2023). Sleep and vigilance states: Embracing spatiotemporal dynamics. Neuron, 111(13), 1998–2011. 10.1016/j.neuron.2023.04.012

Podvalny, E., King, L. E., & He, B. J. (2021). Spectral signature and behavioral consequence of spontaneous shifts of pupil-linked arousal in human. eLife, 10, e68265. 10.7554/eLife.68265

Rattenborg, N. C., Amlaner, C. J., & Lima, S. L. (2000). Behavioral, neurophysiological and evolutionary perspectives on unihemispheric sleep. Neuroscience & Biobehavioral Reviews, 24(8), 817–842. 10.1016/S0149-7634(00)00039-7

Reicher, V., Kis, A., Simor, P., Bódizs, R., & Gácsi, M. (2021). Interhemispheric asymmetry during NREM sleep in the dog. Scientific Reports, 11(1), 18817. 10.1038/s41598-021-98178-3

Reimer, J., Froudarakis, E., Cadwell, C. R., Yatsenko, D., Denfield, G. H., & Tolias, A. S. (2014). Pupil Fluctuations Track Fast Switching of Cortical States during Quiet Wakefulness. Neuron, 84(2), 355–362. 10.1016/j.neuron.2014.09.033

Schneider, M., Hathway, P., Leuchs, L., Sämann, P. G., Czisch, M., & Spoormaker, V. I. (2016). Spontaneous pupil dilations during the resting state are associated with activation of the salience network. NeuroImage, 139, 189–201. 10.1016/j.neuroimage.2016.06.011

Schwarz, L. A., & Luo, L. (2015). Organization of the Locus Coeruleus-Norepinephrine System. Current Biology, 25(21), R1051–R1056. 10.1016/j.cub.2015.09.039

Shine, J. M. (2019). Neuromodulatory Influences on Integration and Segregation in the Brain. Trends in Cognitive Sciences, 23(7), 572–583. 10.1016/j.tics.2019.04.002

Shine, J. M., Koyejo, O., Bell, P. T., Gorgolewski, K. J., Gilat, M., & Poldrack, R. A. (2015). Estimation of dynamic functional connectivity using Multiplication of Temporal Derivatives. NeuroImage, 122, 399–407. 10.1016/j.neuroimage.2015.07.064

Shine, J. M., Lewis, L. D., Garrett, D. D., & Hwang, K. (2023). The impact of the human thalamus on brain-wide information processing. Nature Reviews Neuroscience, 24(7), 416–430. 10.1038/s41583-023-00701-0

Shulman, G. L., Pope, D. L. W., Astafiev, S. V., McAvoy, M. P., Snyder, A. Z., & Corbetta, M. (2010). Right Hemisphere Dominance during Spatial Selective Attention and Target Detection Occurs Outside the Dorsal Frontoparietal Network. The Journal of Neuroscience, 30(10), 3640–3651. 10.1523/JNEUROSCI.4085-09.2010

Siclari, F., & Tononi, G. (2017). Local aspects of sleep and wakefulness. Current Opinion in Neurobiology, 44, 222–227. 10.1016/j.conb.2017.05.008

Sobczak, F., Pais-Roldán, P., Takahashi, K., & Yu, X. (2021). Decoding the brain state-dependent relationship between pupil dynamics and resting state fMRI signal fluctuation. eLife, 10, e68980. 10.7554/eLife.68980

Sturm, W., & Willmes, K. (2001). On the Functional Neuroanatomy of Intrinsic and Phasic Alertness. NeuroImage, 14(1), S76–S84. 10.1006/nimg.2001.0839

Tagliazucchi, E., Roseman, L., Kaelen, M., Orban, C., Muthukumaraswamy, S. D., Murphy, K., Laufs, H., Leech, R., McGonigle, J., Crossley, N., Bullmore, E., Williams, T., Bolstridge, M., Feilding, A., Nutt, D. J., & Carhart-Harris, R. (2016). Increased Global Functional Connectivity Correlates with LSD-Induced Ego Dissolution. Current Biology, 26(8), 1043–1050. 10.1016/j.cub.2016.02.010

Tamaki, M., Bang, J. W., Watanabe, T., & Sasaki, Y. (2016). Night Watch in One Brain Hemisphere during Sleep Associated with the First-Night Effect in Humans. Current Biology, 26(9), 1190–1194. 10.1016/j.cub.2016.02.063

Tanner, J. C., Faskowitz, J., Byrge, L., Kennedy, D. P., Sporns, O., & Betzel, R. F. (2023). Synchronous high-amplitude co-fluctuations of functional brain networks during movie-watching. Imaging Neuroscience, 1, imag–1–00026. 10.1162/imag_a_00026

Thomas Yeo, B. T., Krienen, F. M., Sepulcre, J., Sabuncu, M. R., Lashkari, D., Hollinshead, M., Roffman, J. L., Smoller, J. W., Zöllei, L., Polimeni, J. R., Fischl, B., Liu, H., & Buckner, R. L. (2011). The organization of the human cerebral cortex estimated by intrinsic functional connectivity. Journal of Neurophysiology, 106(3), 1125–1165. (21653723). 10.1152/jn.00338.2011

Van Essen, D. C., Smith, S. M., Barch, D. M., Behrens, T. E. J., Yacoub, E., Ugurbil, K., & W. U-Minn HCP Consortium. (2013). The WU-Minn Human Connectome Project: An overview. NeuroImage, 80, 62–79. 10.1016/j.neuroimage.2013.05.041

Weijs, M. L., Missura, S., Potok-Szybińska, W., Bächinger, M., Badii, B., Carro-Domínguez, M., Wenderoth, N., & Meissner, S. N. (2025). Modulating cortical excitability and cortical arousal by pupil self-regulation. Nature Communications, 16(1), 4552. 10.1038/s41467-025-59837-5

Yang, H., Wu, G., Li, Y., Xu, X., Cong, J., Xu, H., Ma, Y., Li, Y., Chen, R., Pines, A., Xu, T., Sydnor, V. J., Satterthwaite, T. D., & Cui, Z. (2025). Connectional axis of individual functional variability: Patterns, structural correlates, and relevance for development and cognition. Proceedings of the National Academy of Sciences, 122(12), e2420228122. 10.1073/pnas.2420228122

Yellin, D., Berkovich-Ohana, A., & Malach, R. (2015). Coupling between pupil fluctuations and resting-state fMRI uncovers a slow build-up of antagonistic responses in the human cortex. NeuroImage, 106, 414–427. 10.1016/j.neuroimage.2014.11.034

