## supplementary_material for "Arousal modulates functional connectivity through structured and hemispherically asymmetric community architecture during wakefulness"

**Supplement text**

**S1. Robustness analysis of split-half reliability and participant-level resampling**

To ensure that the identified community structures and hemispheric asymmetries were not driven by specific sample compositions or outliers, we performed extensive robustness validations using two complementary resampling strategies: split-half reliability and participant-level resampling. First, we conducted a split-half reliability test with 500 iterations. In each iteration, the total dataset (n = 485 runs) was randomly partitioned into two independent subsets, and the entire analytical pipeline—including community detection and asymmetry quantification—was independently re-executed for each subset. The spatial consistency of community labels across iterations was quantified using the Hungarian algorithm for label alignment and evaluated via Dice coefficients (Figure S1). Second, to account for the potential dependency of multiple sessions within the same participant, we performed a participant-level resampling analysis. We constructed a sub-dataset by randomly selecting only one session per participant (N = 139 participants) over 500 iterations.

The stability of asymmetry results was evaluated across multiple biological scales and analytical dimensions to capture the full landscape of the arousal-modulated connectome. This multi-level assessment encompassed: (i) network-pair hemispheric asymmetry (Figure S2), (ii) regional affiliation patterns (Figure S3), and (iii) community-averaged arousal-tvFC coupling (Figure S4). For each metric, we quantified stability by calculating the absolute deviation (Δ) between the discovery sample and the resampled mean, alongside the relative error percentage. Crucially, when evaluating relative changes, we implemented a thresholding strategy to prevent the statistical inflation of relative error in regions with low baseline effects. In regions or network pairs where the original hemispheric bias was near zero (<0.05), even trivial absolute fluctuations could produce misleadingly large relative errors. By explicitly omitting these inflated numerical annotations, we focused our robustness assessment on functionally relevant and robustly lateralized patterns. This granular approach demonstrates that the reported network-pair specific asymmetries and regional biases are consistent across diverse data partitions, reflecting an intrinsic and stable neurobiological principle of the human brain.

**S2. Sensitivity analysis of sliding-window parameters and temporal lags**

To evaluate the robustness of the identified community structures and their hemispheric asymmetries, we performed extensive sensitivity analyses across varying methodological parameters.

First, we assessed the dependence of our results on sliding-window configurations. The analytical pipeline was repeated using a range of window lengths (30 s, 35 s, 60 s, and 90 s) and sliding steps (1 s, 5 s, and 10 s). The spatial similarity between the resulting community topologies and the original configuration (window length = 30 s, step = 5 s) was quantified using Dice coefficients. As shown in Figure S5, the Dice coefficients for both the overall community structure and individual communities remained consistently above 0.8 across various window parameters (Figure S5A, B, D). These findings demonstrate that the observed arousal-modulated connectivity organization is not dependent on specific sliding-window settings.

Second, considering the hemodynamic response function (HRF) delay and the complex temporal coupling between pupil fluctuations and neural activity, we conducted a lagged cross-correlation analysis. Following established literature on the temporal relationship between tvFC and pupil dynamics, we introduced temporal lags ranging from -3 TR to +3 TR (-3s to 3s) to the pupil time course. We then re-estimated the community structures and assessed their consistency using Dice coefficients. The results (Figure S5C) show that the core community partitions, and lateralization patterns remain stable within physiologically plausible temporal lags, confirming that our primary findings are not biased by the zero-lag assumption.

**S3. Robustness analysis of controlling for nuisance covariates**

To ensure that the identified community structures and their hemispheric asymmetries were not confounded by head motion or non-specific physiological noise, we performed an additional robustness analysis by including nuisance covariates. This is particularly important as pupil diameter, our primary arousal index, is known to covary with head motion, blinks, and non-specific physiological noise, which could potentially bias the estimation of arousal-tvFC coupling.

Specifically, when calculating the arousal-tvFC coupling, we employed partial correlation to regress out the following variables: (i) head motion parameters (quantified by Framewise Displacement, FD) and (ii) the global signal (defined as the mean signal across all gray matter voxels). We then re-executed the community detection using these controlled coupling maps.

As shown in Figure S6, the resulting community topologies remained highly consistent with our primary findings (Figure S6A-C), with Dice coefficients remaining stable across various lag settings even after accounting for these noise components. It is important to note that we did not regress out the eye close ratio, as cumulative evidence suggests that eyelid closure is an intrinsic and valid indicator of arousal levels; controlling for it would likely diminish the biologically relevant coupling between arousal fluctuations and functional connectivity (Sommer & Golz, 2010; Chang et al., 2016; Gonzalez-Castillo et al., 2022). These results confirm that the reported arousal-modulated connectivity organization reflects genuine neural dynamics rather than motion or global artifacts.

**S4 Control analysis on the Optimal Cluster Number**

To ensure that the identified connectivity communities reflect a stable and intrinsic organizational feature of the brain rather than sensitivity to specific methodological decisions, we expanded our evaluation of the clustering solutions using multiple complementary criteria.

As shown in Figure S7, we evaluated the clustering performance for K ranging from 2 to 15 using four distinct metrics: (i) Within-Cluster Sum of Squares (WCSS), reflecting cluster compactness; (ii) Davies-Bouldin Index (DBI), assessing the ratio of within-cluster distances to between-cluster distances; (iii) Calinski-Harabasz (CH) Score, measuring the ratio of between-cluster dispersion to within-cluster dispersion; and (iv) Silhouette Coefficient, evaluating how similar an object is to its own cluster compared to other clusters (Figure S7A).

To objectively identify the "elbow" point where increasing K yields diminishing returns, we applied the L-method (Salvador & Chan, 2004) to the evaluation curves. Specifically, we minimized the total root mean squared error (RMSE) of a two-line linear regression fit across the range of K. The RMSE reached its minimum at K=7 (Figure S7B), represents the optimal balance between model complexity and data fit. Furthermore, we visualized the community topologies across different K values (Figure S7C) to demonstrate that the core spatial organizational principles (e.g., the separation of unimodal and transmodal connectivity) remain consistent across a reasonable range of K. These multi-dimensional evaluations collectively confirm that seven communities provide a parsimonious and stable representation of the arousal-modulated connectome.

**S5. Robustness of pupil diameter time courses across different preprocessing pipelines**

Given that pupil diameter serves as the primary index of arousal in this study, it is critical to ensure that the estimated arousal fluctuations are not idiosyncratic to specific preprocessing choices. We systematically evaluated the robustness of the pupil diameter time courses by comparing our original pipeline with 18 alternative preprocessing combinations.

These pipelines involved variations in three key parameters: (i) temporal smoothing (window sizes of 100 ms, 200 ms, and 500 ms), (ii) artifact interpolation (linear vs. cubic spline), and (iii) blink buffering (durations of 25 ms, 50 ms, and 100 ms). For each fMRI run, we calculated the Pearson correlation coefficient between the pupil time courses derived from these alternative pipelines and the original one.

As illustrated in Figure S8, the mean correlation coefficients across all runs remained exceptionally high (all r > 0.65) regardless of the parameter combinations. The minimal variance (indicated by SD) further confirms the stability of the pupil signal across different sessions and participants. These results demonstrate that the extracted arousal dynamics are highly robust to reasonable changes in pupillometry preprocessing rules, ensuring the reliability of the subsequent arousal-tvFC coupling analyses.

**S6. Data Quality Control and Sample Retention**

To ensure the reliability of the arousal-connectivity analysis, we implemented a rigorous quality control (QC) pipeline for the fMRI data. The participant-level data retention and the distribution of valid runs are summarized in Figure S9. Out of the initial dataset, a total of 139 participants (contributing 485 runs) met the inclusion criteria, which required at least one high-quality resting-state session per subject.

The distribution of available data was characterized across two dimensions: (i) Run-wise consistency per participant, showing the number of participants who contributed one to four valid runs (Figure S9A), and (ii) Session-specific availability, mapping the retention of specific runs across the four sessions (REST1 to REST4) for each individual subject (Figure S9B). This detailed mapping ensures that the longitudinal sampling density was sufficient to capture stable individual arousal-connectivity coupling while accounting for potential data loss due to excessive head motion or physiological artifacts.

**Supplement figure**

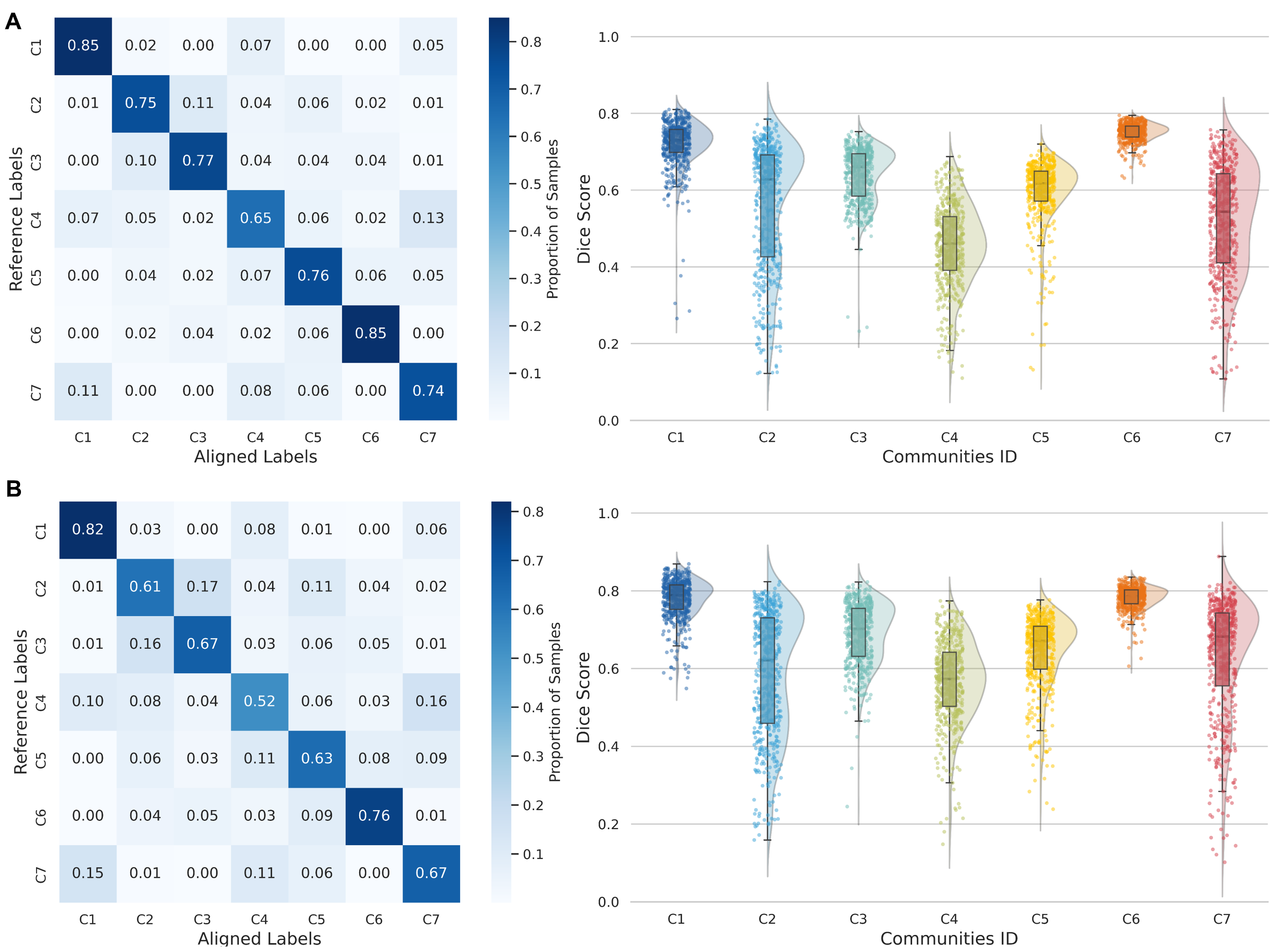

**Figure S1. Stability of community architecture across different resampling strategies. (A)** Confusion matrix (left) showing the consistency between community labels from 500 iterations (aligned via the Hungarian algorithm) and the primary results. The right panel displays the distribution of Dice coefficients for each individual community label. **(B)** Stability of community architecture using participant-level resampling, following the same analytical framework and visualization as in (A).

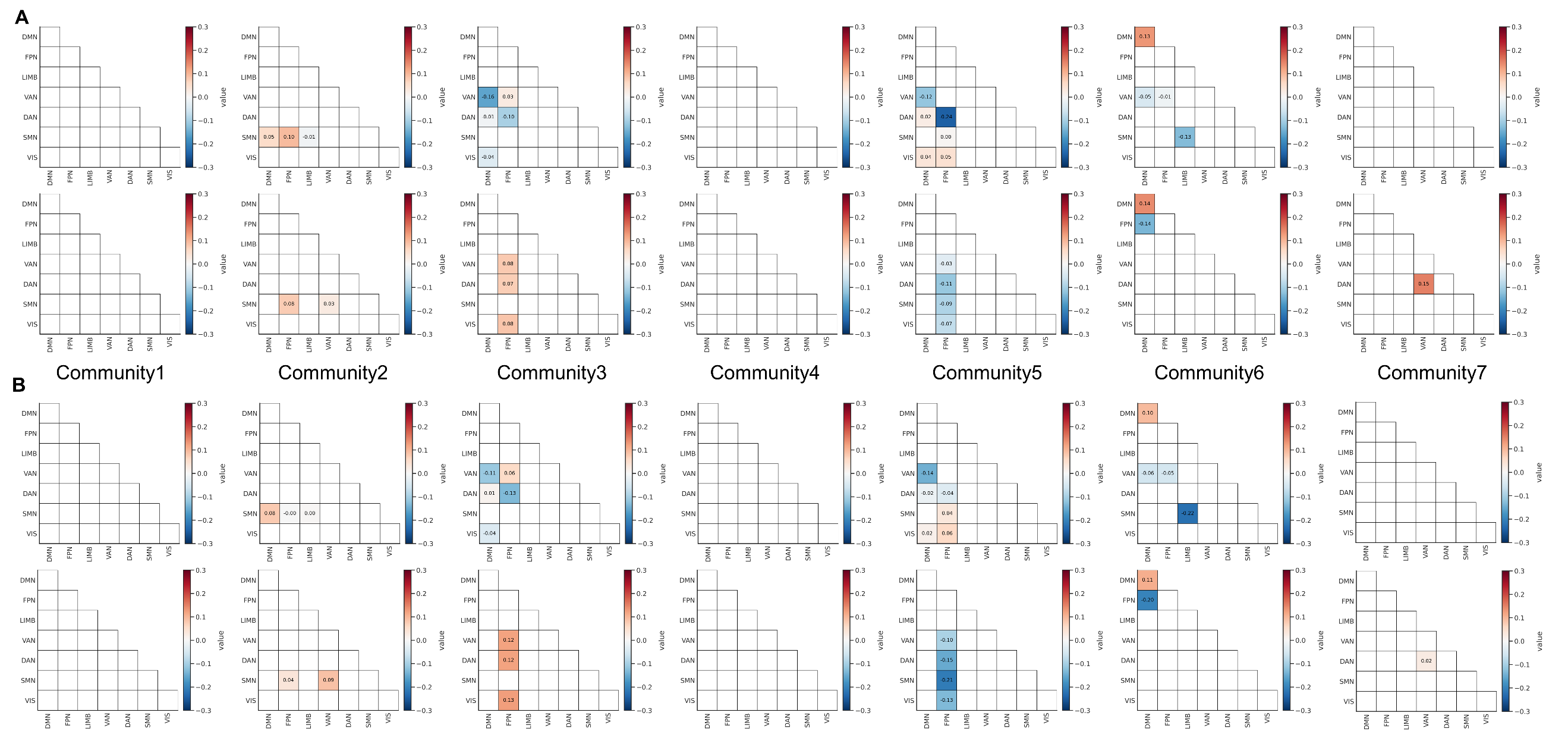

**Figure S2. Robustness of network-pair specific hemispheric asymmetry in community architecture across resampling strategies.** **(A)** The relative value was calculated as: ${{(LI}_{iter} - {LI}_{orig})}/{{LI}_{orig}}$, where ${LI}_{iter}$ represents the mean of 500 resampling iterations and，${LI}_{orig}$ denotes the original findings. The top and bottom rows represent integration and segregation patterns, respectively. **(B)** Stability assessment using participant-level resampling, following the same calculation and visualization framework as in (A).

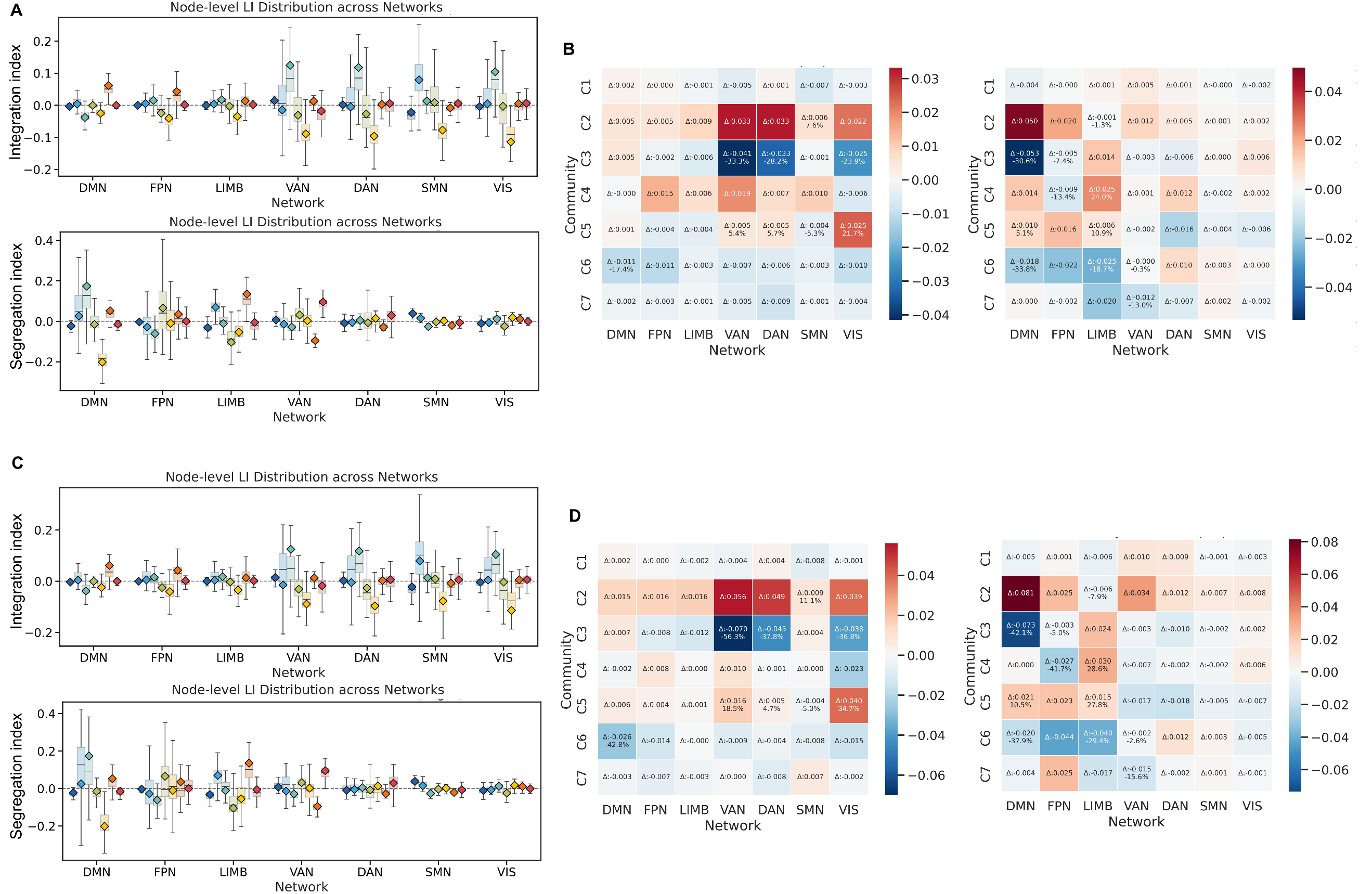

**Figure S3. Robustness of regional affiliation asymmetry across resampling strategies.** **(A)** Stability analysis using split-half resampling. Boxplots represent the distribution of asymmetry indices across 500 iterations, where colored diamonds indicate the original findings. **(B)** The heatmap background represents the absolute deviation (Δ) between the discovery sample and the resampled mean. Numerical annotations indicate the relative error percentage (%). To prevent the statistical inflation of relative values in regions with minimal effects, these annotations are explicitly omitted where the ${LI}_{orig}$ is low (< 0.05), focusing the assessment on robustly lateralized regions. **(C-D)** Stability assessment using participant-level resampling, following the same quantitative framework. The overall tight alignment between resampled distributions and original discovery values confirms that the spatial patterns of regional hemispheric bias are highly stable and not artifacts of specific sample selection.

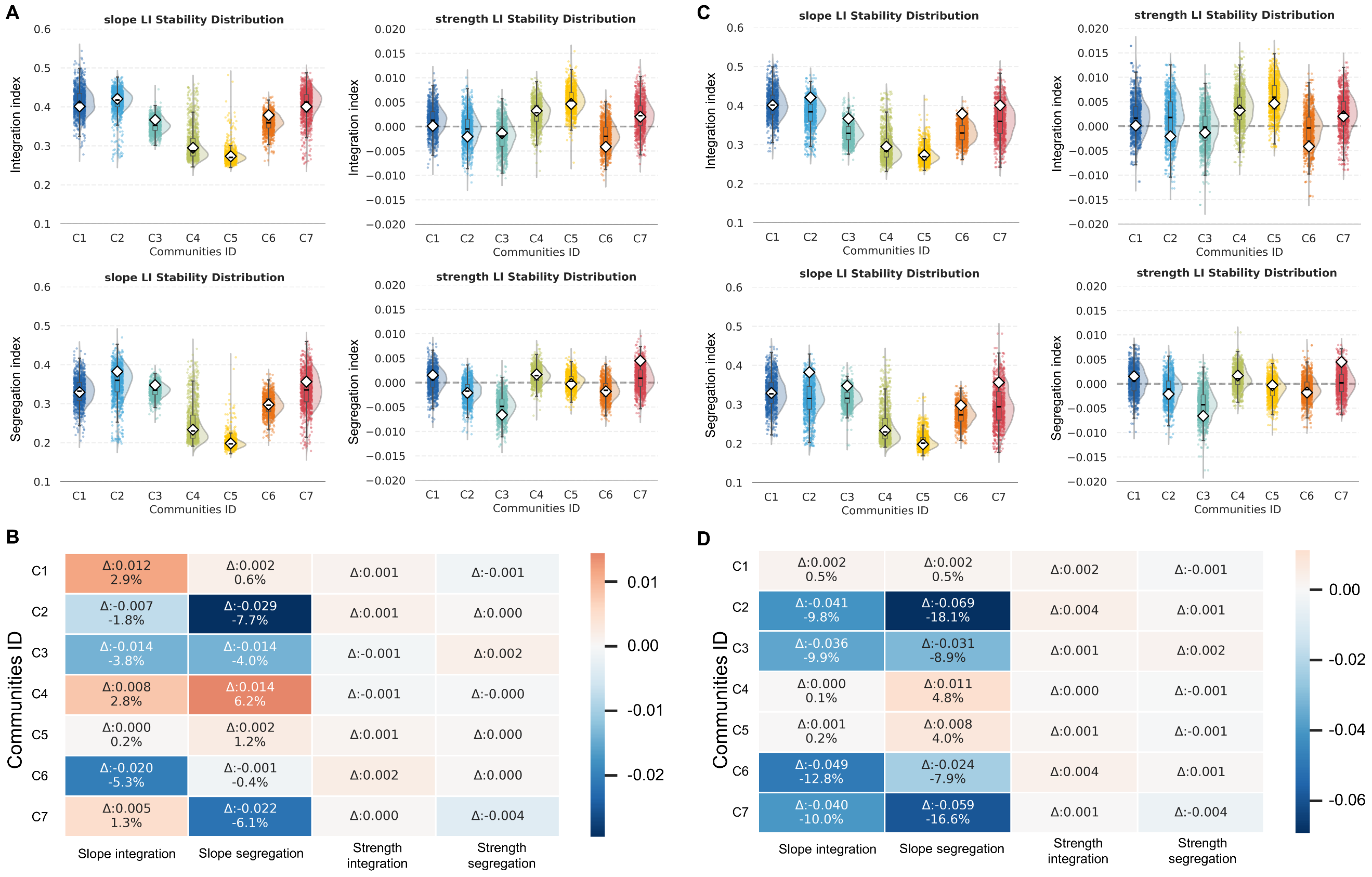

**Figure S4. Reliability of arousal-FC coupling asymmetry across resampling strategies. (A)** Reliability assessment using split-half resampling. Boxplots illustrate the distribution of asymmetry indices for arousal-FC coupling across 500 iterations, with diamonds indicating the original discovery findings. (B) The heatmap background represents the absolute deviation (Δ) between the discovery sample and the resampled mean. Numerical annotations indicate the relative error percentage (%). To prevent the statistical inflation of relative values in network pairs with minimal effects, these annotations are explicitly omitted where the ${LI}_{orig}$ is low (< 0.05), focusing the assessment on robustly lateralized patterns. **(C-D)** Reliability assessment using participant-level resampling, following the same quantitative framework. The consistent alignment between the resampled distributions and primary findings confirms that the hemispheric asymmetry of arousal-tvFC coupling is a robust biological feature rather than a result of sampling bias or specific data partitions.

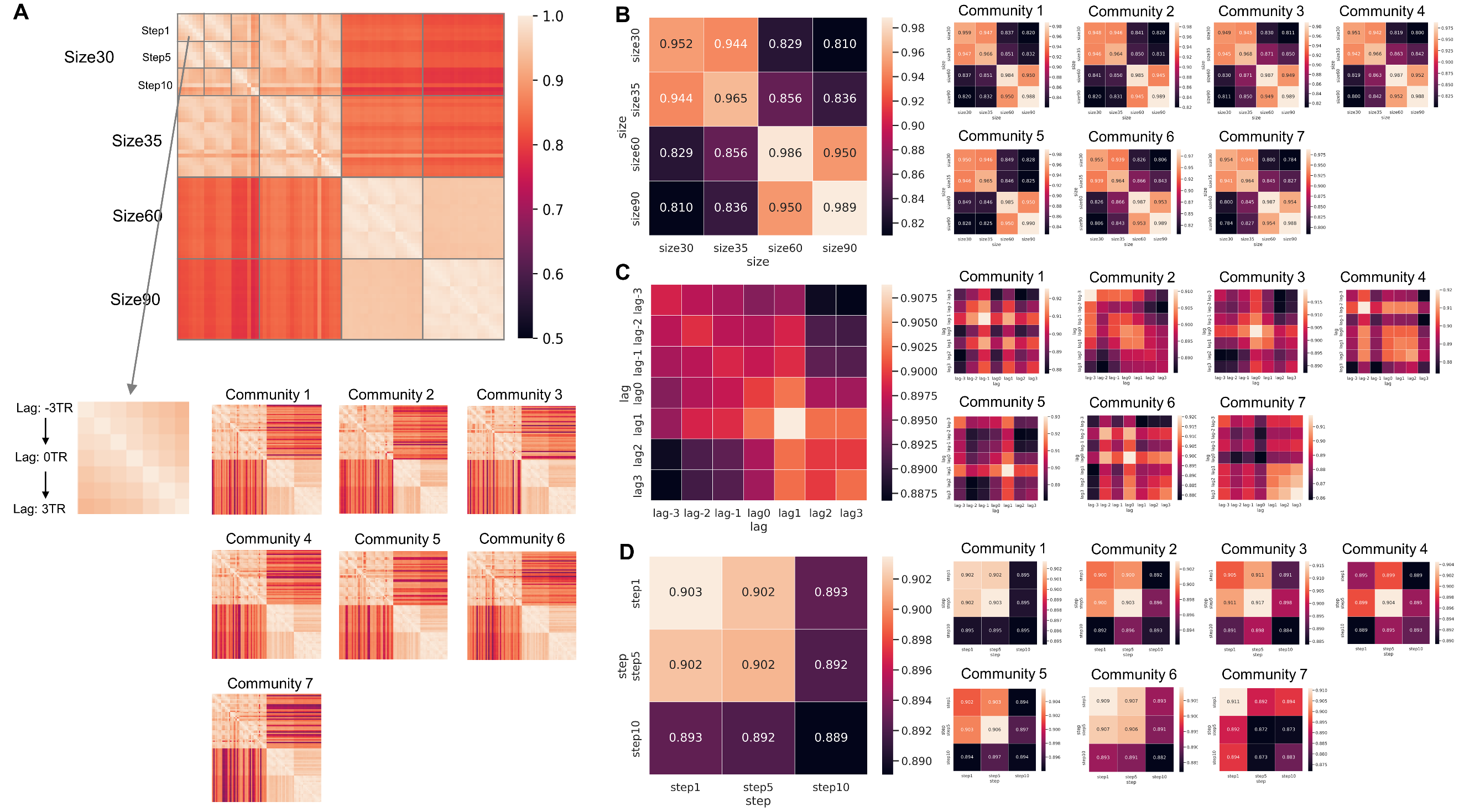

**Figure S5. Robustness of community detection across sliding-window parameters and temporal lags. (A)** Dice coefficients between the community topologies derived from different combinations of window size, window step, and temporal lag. The top heatmap illustrates the Dice coefficients for the overall community partition, while the bottom heatmaps show the coefficients calculated independently for each individual community. **(B–D)** Marginal consistency matrices for specific parameters. These panels display the Dice coefficients averaged across all other parameters while retaining only (B) window size, (C) temporal lag, and (D) window step.

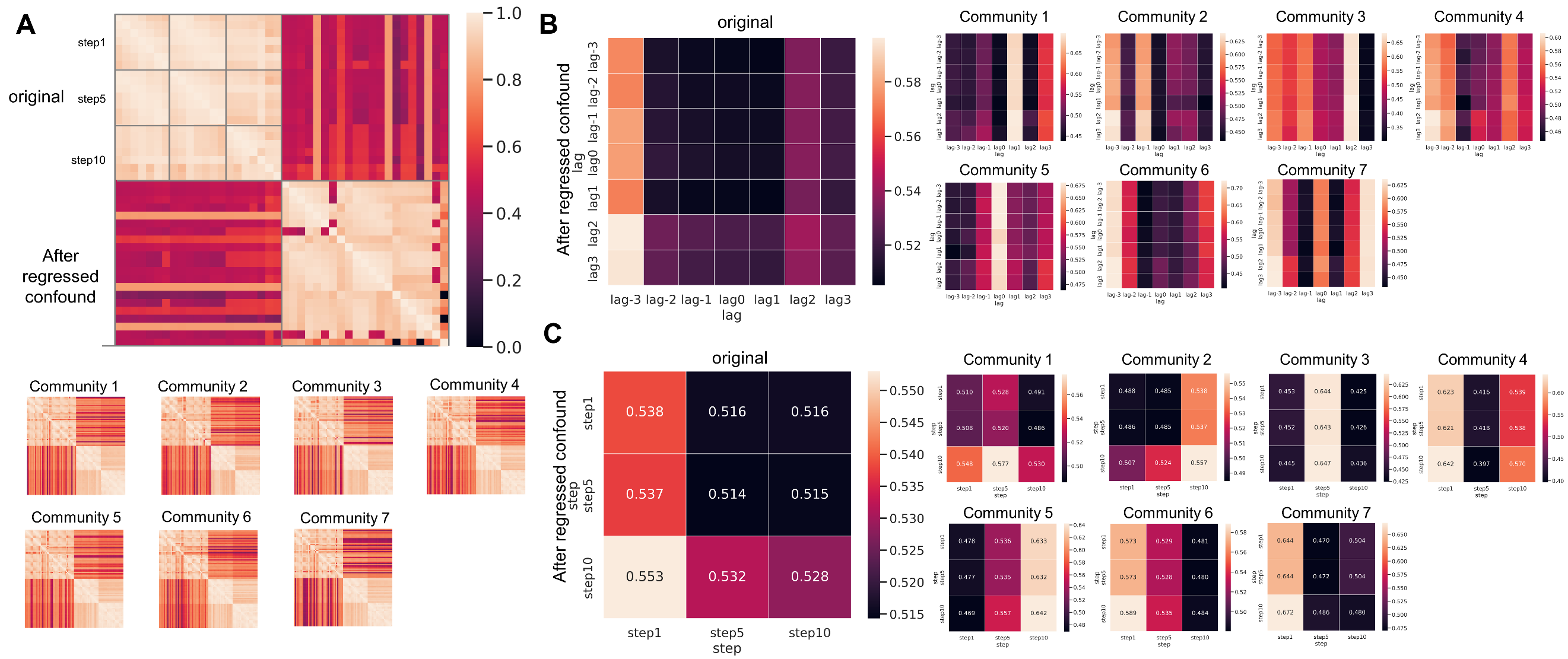

**Figure S6. Robustness of community architecture after controlling for motion and global signal artifacts. (A)** Dice coefficients between the original community templates and those derived after regressing out nuisance covariates (Framewise Displacement and Global Signal) across various window steps and temporal lags. The top heatmap shows the Dice coefficients for the overall community partition, while the bottom heatmaps display the coefficients for each individual community. **(B)** Marginal Dice coefficient matrix averaged across all parameters except for the temporal lag. **(C)** Marginal Dice coefficient matrix averaged across all parameters except for the window step.

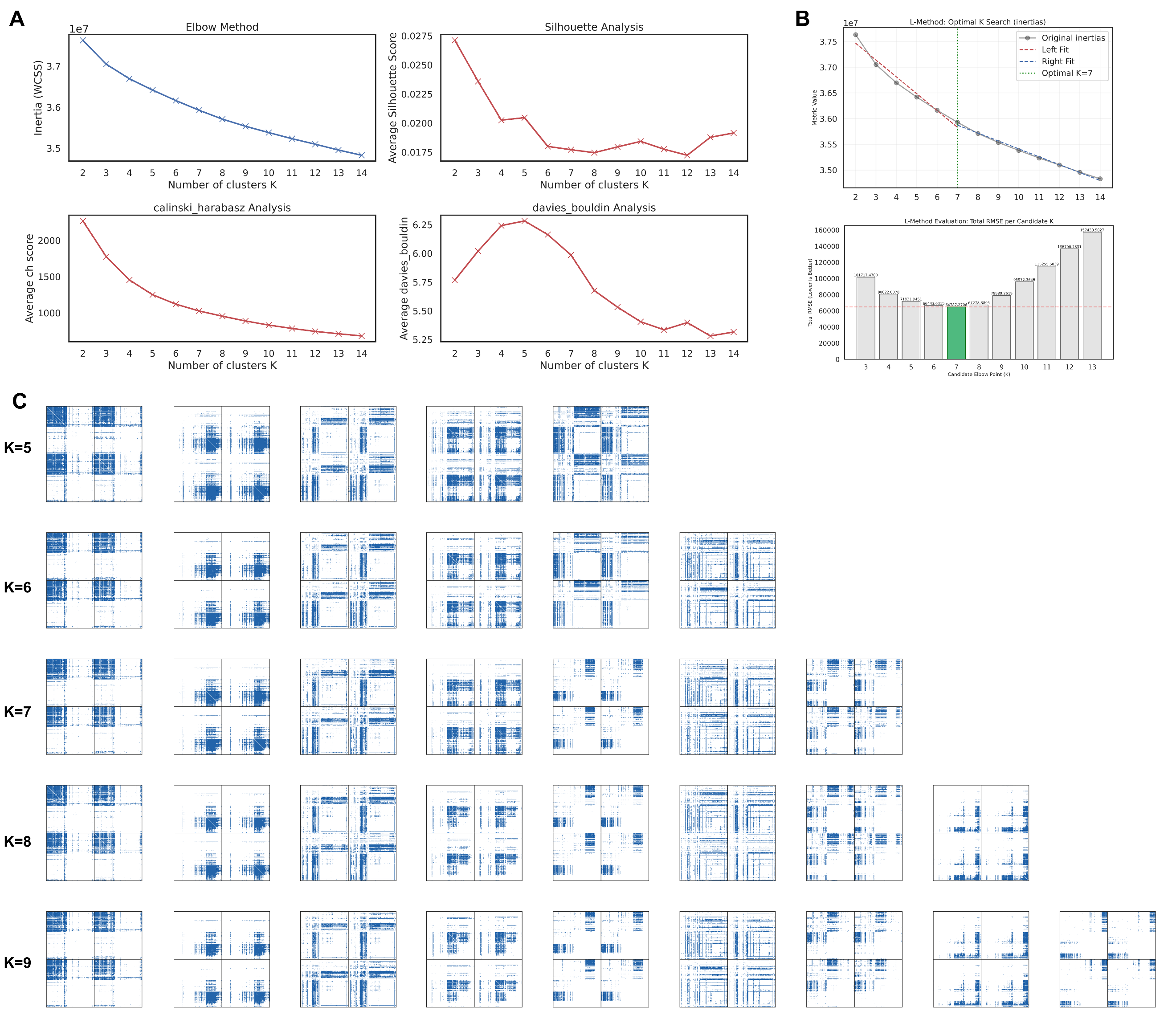

**Figure S7. Evaluation and selection of the optimal number of communities. (A)** Evaluation curves for four complementary clustering quality metrics across K=2 to 15: Within-Cluster Sum of Squares (WCSS), Davies-Bouldin Index, Calinski-Harabasz Score, and Silhouette Coefficient. **(B)** Objective determination of the "knee" point using the L-method. The plot shows the total root mean squared error (RMSE) of the two-line linear regression fit as a function of K, with the minimum RMSE achieved at K=7 (indicated by the arrow). **(C)** Spatial distribution of community labels across k=5 to 9. The visualization demonstrates the hierarchical evolution and spatial stability of the community topologies as the number of clusters increases.

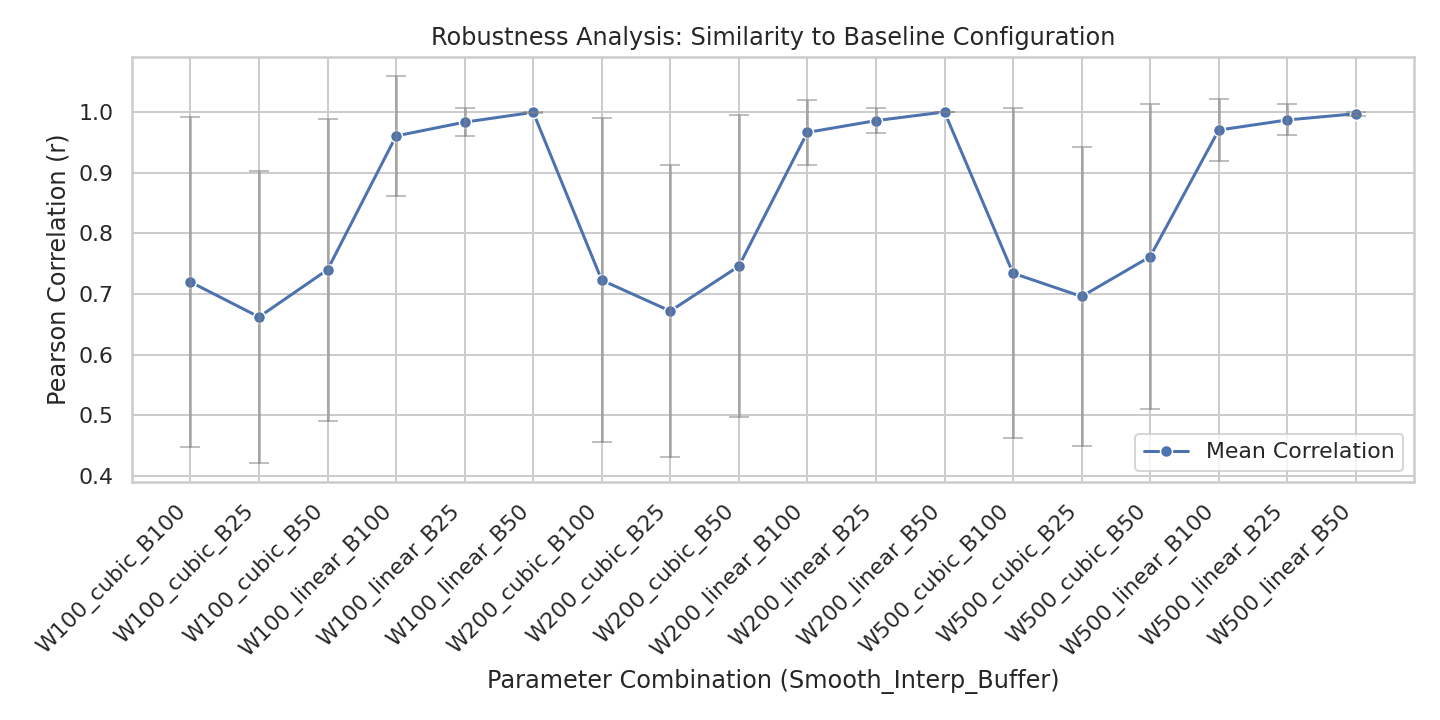

**Figure S8. Robustness of pupil diameter time courses across different preprocessing pipelines.** Line plots illustrate the Pearson correlation coefficients between the pupil diameter time courses derived from the original pipeline and those from 18 alternative preprocessing combinations. These pipelines varied across three parameters: (i) Smoothing window size (100 ms, 200 ms, and 500 ms); (ii) Interpolation method (linear or cubic spline); and (iii) Blink buffer duration (25 ms, 50 ms, and 100 ms). Each data point represents the mean correlation across all fMRI runs, with error bars indicating ±1 standard deviation (SD).

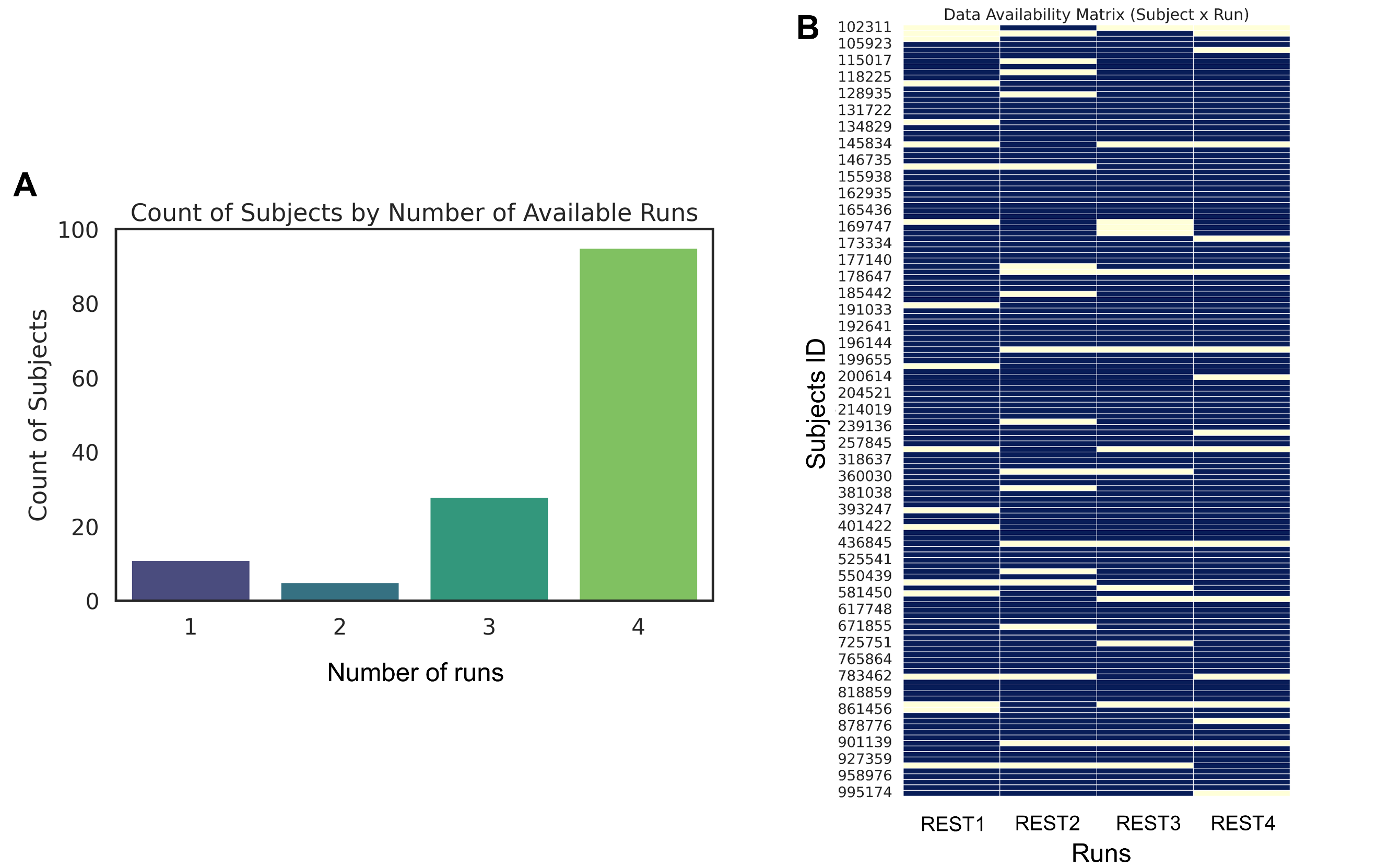

**Figure S9. Participant-level data retention and run distribution after quality control.** **(A)** Summary of valid runs per participant. The bar chart shows the number of participants who contributed 1, 2, 3, or 4 fMRI runs to the final analysis (N = 139 participants). **(B)** Individual run availability matrix. The heatmap illustrates the distribution of retained runs across the four sessions (REST1 to REST4) for each participant. Each row represents an individual participant, and columns represent the four scanning sessions. Colored cells indicate that the specific run passed all quality control (QC) criteria, while white cells indicate excluded or unavailable runs.

**Reference**

Chang, C., Leopold, D. A., Schölvinck, M. L., Mandelkow, H., Picchioni, D., Liu, X., Ye, F. Q., Turchi, J. N., & Duyn, J. H. (2016). Tracking brain arousal fluctuations with fMRI. *Proceedings of the National Academy of Sciences*, *113*(16), 4518–4523. https://doi.org/10/f8ktgg

Gonzalez-Castillo, J., Fernandez, I. S., Handwerker, D. A., & Bandettini, P. A. (2022). Ultra-slow fMRI fluctuations in the fourth ventricle as a marker of drowsiness. *NeuroImage*, *259*, 119424. https://doi.org/10.1016/j.neuroimage.2022.119424

Salvador, S., & Chan, P. (2004). Determining the number of clusters/segments in hierarchical clustering/segmentation algorithms. *16th IEEE International Conference on Tools with Artificial Intelligence*, 576–584. https://doi.org/10.1109/ICTAI.2004.50

Sommer, D., & Golz, M. (2010). Evaluation of PERCLOS based current fatigue monitoring technologies. *2010 Annual International Conference of the IEEE Engineering in Medicine and Biology*, 4456–4459. https://doi.org/10.1109/IEMBS.2010.5625960
